# Meaningful Associations in the Adolescent Brain Cognitive Development Study

**DOI:** 10.1101/2020.09.01.276451

**Authors:** Anthony Steven Dick, Daniel A. Lopez, Ashley L. Watts, Steven Heeringa, Chase Reuter, Hauke Bartsch, Chun Chieh Fan, David N. Kennedy, Clare Palmer, Andrew Marshall, Frank Haist, Samuel Hawes, Thomas E. Nichols, Deanna M. Barch, Terry L. Jernigan, Hugh Garavan, Steven Grant, Vani Pariyadath, Elizabeth Hoffman, Michael Neale, Elizabeth A. Stuart, Martin P. Paulus, Kenneth J. Sher, Wesley K. Thompson

## Abstract

The Adolescent Brain Cognitive Development (ABCD) Study is the largest single-cohort prospective longitudinal study of neurodevelopment and children’s health in the United States. A cohort of n= 11,880 children aged 9-10 years (and their parents/guardians) were recruited across 22 sites and are being followed with in-person visits on an annual basis for at least 10 years. The study approximates the US population on several key sociodemographic variables, including sex, race, ethnicity, household income, and parental education. Data collected include assessments of health, mental health, substance use, culture and environment and neurocognition, as well as geocoded exposures, structural and functional magnetic resonance imaging (MRI), and whole-genome genotyping. Here, we describe the ABCD Study aims and design, as well as issues surrounding estimation of meaningful associations using its data, including population inferences, hypothesis testing, power and precision, control of covariates, interpretation of associations, and recommended best practices for reproducible research, analytical procedures and reporting of results.

## 1.0 Introduction

The Adolescent Brain Cognitive Development^SM^ (ABCD) Study is the largest single-cohort long-term longitudinal study of neurodevelopment and child and adolescent health in the United States. The study was conceived and initiated by the United States’ National Institutes of Health (NIH), with funding beginning on September 30, 2015. The ABCD Study^®^ collects observational data to characterize US population trait distributions and to assess how biological, psychological, and environmental factors (including interpersonal, institutional, cultural, and physical environments) can influence how individuals live and develop in today’s society. From the outset, the NIH and ABCD scientific investigators were motivated to develop a baseline sample that reflected the sociodemographic variation present in the US population of 9-10 year-old children, and to follow them longitudinally through adolescence and into early adulthood.

Population representativeness, or more precisely, absence of uncorrected selection bias in the subject pool, is important in achieving external validity, i.e., the ability to generalize specific results of the study to US society at large. As described below, the ABCD Study attempted to match the diverse US population of 9-10 year-old children on key demographic characteristics. However, even with a largely representative sample, failure to account for key covariates can affect internal validity, i.e., the degree to which observed associations accurately reflect the effects of underlying causal mechanisms. Moreover, it is crucial that the study collects a rich array of variables that may act as moderators or mediators, including biological and environmental variables, in order to aid in identifying potentially causal pathways of interest, to quantify individualized risk for (or resilience to) poor outcomes, and to inform public policy decisions. External and internal validity also depend on assessing the impact of random and systematic measurement error, implementing analytical methods that incorporate relevant aspects of study design, and emphasizing robust and replicable estimation of associations.

The ABCD cohort is large enough that very small effects related to developmental outcomes can be reliably estimated for many developmental outcomes. It is therefore directly addressing the over-estimation of association sizes and replication issues affecting much of current behavioral and neuroscience research^1,2^. Given the large sample size of the ABCD cohort, emphasis should be placed on accurate and replicable estimation of associations rather than mere statistical significance. Indeed, a primary strength of the ABCD Study is that more accurate assessment of the magnitude of associations promotes realistic judgments as to their relevance and utility for understanding mechanisms, for precision medicine, and for public health policy.

Furthermore, a large sample size and rich assessment protocol enable the construction of more realistically complex etiological models which simultaneously incorporate factors from multiple domains. Even if the effects of individual factors are small, as has been the case in other large epidemiological samples^3,4^, they may still be useful for uncovering the genetic and environmental mechanisms of neurodevelopment, behavior, and health. Observed associations may be small (e.g., due to measurement error) even if the underlying effects are biologically important^5^. Moreover, many small biological effects may in concert explain a sizeable proportion of the variation in neurodevelopmental trajectories, as has been recently demonstrated in genome-wide association analyses of complex traits^6^. Effects may also accumulate or become larger as subjects pass through adolescence into early adulthood^7^.

The ABCD Study was conceived to address some of the most important public health questions facing today’s children and adolescents^8^. These questions include identifying factors leading to the initiation and consumption patterns of psychoactive substances, substance-related problems, and substance use disorders as well as their subsequent impact on the brain, neurocognition, health, and mental health over the course of adolescence and into early adulthood. More broadly, a large epidemiologically informed longitudinal study beginning in childhood and continuing on through early adulthood will provide a wealth of unique data on normative development, as well as environmental and biological factors associated with variation in developmental trajectories. This broader perspective has led to the involvement of multiple NIH Institutes that are stakeholders in the range of health outcomes targeted in the ABCD design. (Information regarding funding agencies, recruitment sites, investigators, and project organization can be obtained at https://abcdstudy.org).

The ABCD Study primary aims are given in the Supplementary Materials (SM) Section S.1. Briefly, these include development of national standards for normal brain development, estimation of individual developmental trajectories of mental and physical health and substance use and their inter-relationships, and assessment of the genetic and environmental factors impacting these trajectories. We describe the study design and outline analytic strategies to address the primary study aims, including worked examples, with emphasis on approaches that incorporate relevant aspects of study design. We emphasize the impact of sample size on the precision of association estimates and thoughtful control of covariates in the context of the large-scale population neuroscience data produced by the ABCD Study. Finally, in the Supplementary Materials we describe state-of-the-field recommendations for promoting reproducible science and briefly outline best practices for statistical analyses and reporting of results using the ABCD Study data.

## 2.0 Study Design

The ABCD Study is a prospective longitudinal cohort study of US children born between 2006-2008. A total cohort of *n* = 11880 children aged 9-10 years at baseline (and their parents/guardians) was recruited from 22 sites (with one site no longer active) and are being followed for at least ten years. Eligible children were recruited from the household populations in defined catchment areas for each of the study sites during the roughly two-year period beginning September 2016 and ending in October 2018.

Within study sites, consenting parents and assenting children were primarily recruited through a probability sample of public and private schools augmented to a smaller extent by special recruitment through summer camp programs and community volunteers. ABCD employed a probability sampling strategy to identify schools within the catchment areas as the primary method for contacting and recruiting eligible children and their parents. This method has been used in other large national studies (e.g., Monitoring the Future^9^; the Add Health Study^10^; the National Comorbidity Replication-Adolescent Supplement^11^; and the National Education Longitudinal Studies^12^). Twins were recruited from birth registries (see^13,14^ for participant recruitment details). A minority of participants were recruited through non-school-based community outreach and word-of-mouth referrals.

Across recruitment sites, inclusion criteria consisted of being in the required age range and able to provide informed consent (parents) and assent (child). Exclusions were minimal and were limited to lack of English language proficiency in the children, the presence of severe sensory, intellectual, medical or neurological issues that would impact the validity of collected data or the child’s ability to comply with the protocol, and contraindications to MRI scanning^13^. Parents must be fluent in either English or Spanish.

Measures collected in the ABCD Study include a neurocognitive battery^15,16^, mental and physical health assessments^17^, measures of culture and environment^18^, biospecimens^19^, structural and functional brain imaging^20,21^, geolocation-based environmental exposure data, wearables and mobile technology^22^, and whole genome genotyping^23^. Many of these measures are collected at in-person annual visits, with brain imaging collected at baseline and at every other year going forward. A limited number of assessments are collected in semi-annual telephone interviews between in-person visits. Data are publicly released on an annual basis through the NIMH Data Archive (NDA, https://nda.nih.gov/abcd). Figure 1 graphically displays the measures that have been collected as part of the ABCD NDA 3.0. Release. Figure 2 depicts the planned data collection and release schedule over the initial 10 years of the study.

**Figure 1:**
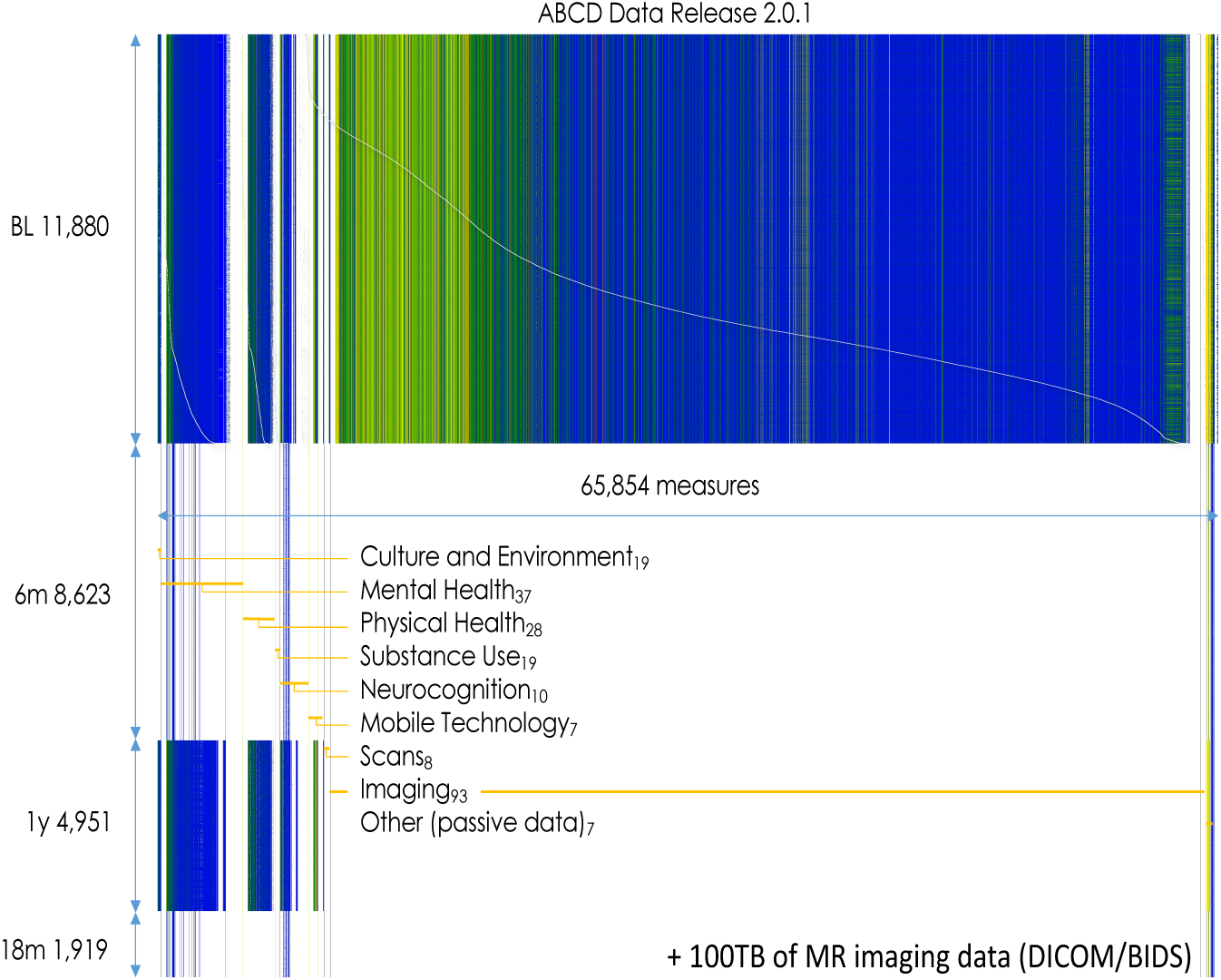
**ABCD Study Assessments for NDA 2.0.1 Release Data**

**Figure 2:**
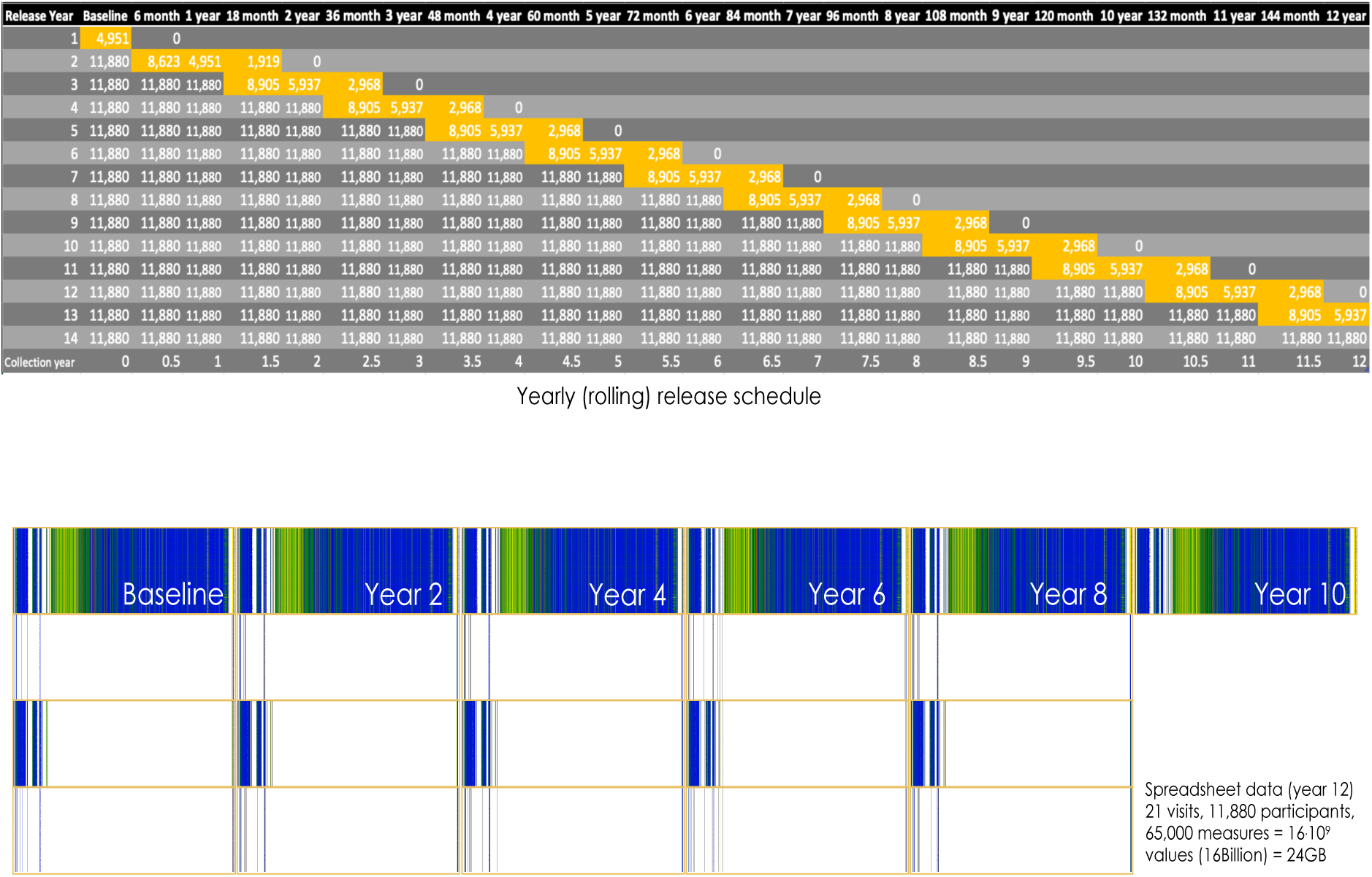
**ABCD Data Collection and NDA Release Schedule**

ABCD sample demographics (from NDA Release 2.0.1, which contains data from *n* = 11875 subjects) are presented in Table 1, along with a comparison to the corresponding statistics from the American Community Survey (ACS). The ACS is a large probability sample survey of US households conducted annually by the US Bureau of Census and provides a benchmark for selected demographic and socio-economic characteristics of US children aged 9-10 years. The 2011-2015 ACS Public Use Microsample (PUMS) file provides data on over 8,000,000 sample US households. Included in this five-year national sample of households are 376,370 individual observations for children aged 9-10 and their households. With some minor differences, the unweighted distributions for the ABCD baseline sample closely match the ACS-based national estimates for demographic characteristics including age, sex, and household size. The general concordance of the samples can be attributed in large part to three factors: 1) the inherent demographic diversity across the ABCD study sites; 2) stratification (by race/ethnicity) in the probability sampling of schools within sites; and 3) demographic controls employed in the recruitment by site teams. Likewise, the unweighted percentages of ABCD children for the most prevalent race/ethnicity categories are an approximate match to the ACS estimates for US children age 9 and 10. Collectively, children of Asian, American Indian/Alaska Native (AIAN) and Native Hawaiian/Pacific Islander (NHPI) ancestry are under-represented in the unweighted ABCD data (3.2%) compared with ACS national estimates (5.9%). This outcome, which primarily affects ABCD’s sample of Asian children, may be due in part to differences in how the parent/caregiver of the child reports multiple race/ethnicity ancestry in ABCD and the ACS.

**Table 1:** ABCD Baseline and ACS 2011-2015 Demographic Characteristics.

| Characteristic | Category | ABCD<br>n=11,879 | ACS 2011-2015 |  |
| --- | --- | --- | --- | --- |
|  |  | % | N | % |
| Population | Total | 100 | 8,211,605 | 100 |
| Age | 9 | 52.3 | 4,074,807 | 49.6 |
|  | 10 | 47.7 | 4,136,798 | 50.4 |
| Sex | Male | 52.2 | 4,205,925 | 51.2 |
|  | Female | 47.8 | 4,005,860 | 48.8 |
| Race/Ethnicity | NH White | 52.2 | 4,305,552 | 52.4 |
|  | NH Black | 15.1 | 1,101,297 | 13.4 |
|  | Hispanic | 20.4 | 1,973,827 | 24.0 |
|  | Asian, AIAN, NHPI | 3.2 | 487,673 | 5.9 |
|  | Multiple | 9.2 | 343,256 | 4.2 |
| Family Income | <\$25k | 16.1 | 1,762,415 | 21.5 |
| | \$25k-\$49k | 15.1 | 1,784,747 | 21.7 |
| | \$50k-\$74k | 14.0 | 1,397,641 | 17.0 |
| | \$75k-\$99k | 14.1 | 1,023,127 | 12.5 |
| | \$100k-\$199k | 29.5 | 1,685,036 | 20.5 |
| | \$200k + | 11.2 | 558,639 | 6.8 |
| Family Type | Married Parents | 73.4 | 5,426,131 | 66.1 |
|  | Other Family Type | 26.6 | 2,785,474 | 33.9 |
| Parent Employment | Married, 2 in LF | 50.2 | 3,353,572 | 40.8 |
|  | Married, 1 in LF | 21.9 | 1,949,288 | 23.7 |
|  | Married, 0 in LF | 1.3 | 156,807 | 1.9 |
|  | Single, in LF | 21.1 | 2,174,365 | 26.5 |
|  | Single, Not in LF | 5.4 | 577,573 | 7.0 |
| Region | Northeast | 16.9 | 1,336,183 | 16.3 |
|  | Midwest | 20.4 | 1,775,723 | 21.6 |
|  | South | 28.3 | 3,117,158 | 38.0 |
|  | West | 34.4 | 1,982,541 | 24.1 |
| Household Size | 2 to 3 | 17.3 | 1,522,216 | 18.5 |
|  | 4 | 33.5 | 2,751,942 | 33.5 |
|  | 5 | 24.9 | 2,085,666 | 25.4 |
|  | 6 | 14.0 | 1,025,285 | 12.5 |
|  | 7+ | 10.3 | 826,496 | 10.1 |

**LF**=labor force

A feature of the ABCD design that deserves attention in the analysis of the baseline cohort data is the special oversample of twin pairs in four of the ABCD sites. Although twins were eligible to be recruited in all sites that used the school-based recruitment sampling methodology, in the four special twin sites supplemental samples of 150-250 twin pairs per site were enrolled in ABCD using twins selected from state registries^13^; . These special samples of twin pairs can be distinguished in the final baseline cohort; however, the study has chosen not to explicitly segregate these twin data from the general population sample of single births and incidental twins recruited through the school-based sampling protocol. The data provide opportunities to assay differences between twins and non-twins, which may potentially limit the generalizability of genetically informed twin analyses to the population as whole.

## 3.0 Population Inferences

The ABCD recruitment effort worked very hard to maintain similarity of the ABCD sample and the US population with respect to sex and race/ethnicity of the children in the study. The predominantly probability sampling methodology for recruiting children within each study site was intended to randomize over confounding factors that were not explicitly controlled (or subsequently reflected in the population weighting). Nevertheless, school consent and parental consent were strong forces that certainly may have altered the effectiveness of the randomization over these uncontrolled confounders.

The purpose of the population weighting described below is to control for specific sources of selection bias and restore unbiasedness to descriptive and analytical estimates of the population characteristics and relationships. For many measures of substantive interest, the success of this effort will never be fully known except in rare cases where comparative national benchmarks exist (e.g. children’s height) from administrative records or very large surveys or population censuses. The first step in benchmarking the ABCD baseline sample weights to population estimates from the ACS sample required identification of a key set of demographic and socio-economic variables for the children and their households that are measured in both the ABCD Study and in the ACS household interviews. For the ABCD eligible children, the common variables include 1) age; 2) sex; and 3) race/ethnicity. For the child’s household, additional variables include: 4) family income; 5) family type (married parents, single parent); 6) household size 7) parents’ work force status (family type by parent employment status); 8) Census Region.

The construction of the population weights is described in detail elsewhere^24^. Briefly, a multiple logistic regression model was fit to the concatenated ACS and ABCD data. In estimating the parameters of this model, each case in the concatenated file receives a frequency weight. ACS cases are assigned their population weights which in aggregate sum to an average estimate of the US population of children age 9, 10 for the period 2011-2015. ABCD cases are assigned a unit weight. Applying the frequency weights in the estimation of the model ensures that the corresponding population propensities for the ABCD sample cases reflect the base population fraction (∼ 0.00145) as well as adjustments for the individual covariate factors in the model. The population weight values for each ABCD case are then obtained by taking the reciprocal of the predicted probability of sample membership for the case, trimming extreme weights, and then “raking” the trimmed initial weights to exact ACS population counts for the marginal categories of age, sex at birth, and race/ethnicity. With case-specific population weights assigned to each subject, weighted estimates and standard errors of population characteristics or parameters in population models can be computed using survey analysis software (such as the survey package^25^ in R) along with robust standard errors and confidence intervals for the weighted estimates^26^. Note, these are weights for the baseline samples; weights reflecting the sample composition at each follow-up will also be developed and disseminated going forward.

Heeringa and Berglund (2020)^24^ present regression analyses with and without using the population weights. Although it is important not to over-generalize from a small set of comparisons to all possible analyses of the ABCD data, the results described therein lead to several recommendations for researchers who are analyzing the ABCD baseline data, summarized below. R scripts for computing the ABCD population weights and for applying them in analyses are available at https://github.com/ABCD-STUDY/abcd_acs_raked_propensity. The population weights computed as described here are available in the NDA data releases 2.0.1 and 3.0.

First, unweighted analysis may result in biased estimates of descriptive population statistics. The potential for bias in unweighted estimates from the ABCD data is strongest when the variable of interest is highly correlated with socio-economic variables including family income, family type and parental work force participation.

Second, for regression models of the ABCD baseline data, an unweighted analysis using mixed-effects models (e.g., site, family, individual) is the preferred choice. Presently, there is no empirical evidence from comparative analyses that methods for multi-level weighting^27^ will improve the accuracy or precision of the model fit, although additional research on this topic is ongoing.

Third, comparative analyses of descriptive estimation methods presented in Heeringa and Berglund (2020)^24^ found that, properly weighted, results for the pooled general population and special twin samples are comparable to those for weighted estimates based solely on the smaller general population sample. Likewise, regression analyses based on the pooled general population and special twin samples that account for inter-familial clustering (e.g., multi-level models) produce similar results to analyses based on the general population sample alone. Nevertheless, analysts should use appropriate caution in pooling the general population and special twin samples for analyses, as the exchangeability observed in the comparative analyses presented in Heeringa and Berglund (2020)^24^ may not necessarily hold in general.

As a demonstration of the implications of the weighting strategy employed in the ABCD Study, weighted and unweighted means and standard errors for ABCD baseline brain morphometry - volumes of cortical Desikan parcels^28^ - are presented in Table 2. Missing observations were first imputed using the R library **mice**^29^ before applying weights to the completed sample. Differences between unweighted and weighted means are quite small in the baseline sample in this case. As longitudinal MRI data become available in ABCD (starting with the second post-baseline annual follow-up visit), population-valid mean trajectories of brain-related outcomes will also be computable using a similar population weighting scheme, also allowing for characterization of variation of trajectories from the population mean.

**Table 2:** Unweighted and Weighted Means of Desikan Cortical Volumes.

|  | Mean | SE | Weighted Mean | SE |
| --- | --- | --- | --- | --- |
| bankssts | 3238.48 | 473.95 | 3227.70 | 472.83 |
| caudalanteriorcingulate | 2571.23 | 476.91 | 2559.34 | 478.06 |
| caudalmiddlefrontal | 8326.70 | 1408.47 | 8277.25 | 1398.77 |
| cuneus | 3645.25 | 582.41 | 3626.44 | 582.07 |
| entorhinal | 1843.15 | 339.44 | 1835.95 | 339.10 |
| fusiform | 12050.11 | 1552.79 | 12009.48 | 1558.06 |
| inferiorparietal | 18387.31 | 2432.67 | 18325.23 | 2428.86 |
| inferiortemporal | 13182.85 | 1879.13 | 13133.08 | 1870.21 |
| isthmuscingulate | 3252.16 | 534.48 | 3239.51 | 538.27 |
| lateraloccipital | 13334.05 | 1870.71 | 13283.90 | 1848.41 |
| lateralorbitofrontal | 9295.28 | 1036.65 | 9258.68 | 1035.60 |
| lingual | 8031.18 | 1132.35 | 7998.54 | 1132.13 |
| medialorbitofrontal | 5976.38 | 731.09 | 5954.65 | 725.41 |
| middletemporal | 14275.50 | 1796.11 | 14230.8 | 1786.83 |
| parahippocampal | 2586.48 | 378.94 | 2576.70 | 378.86 |
| paracentral | 4674.33 | 672.68 | 4660.61 | 674.30 |
| parsopercularis | 5701.08 | 849.03 | 5683.61 | 846.91 |
| parsorbitalis | 3097.73 | 371.12 | 3084.29 | 371.66 |
| parstriangularis | 5178.54 | 733.71 | 5159.42 | 732.41 |
| pericalcarine | 2505.86 | 425.52 | 2489.51 | 424.71 |
| postcentral | 11822.49 | 1599.97 | 11788.14 | 1593.43 |
| posteriorcingulate | 4196.07 | 603.72 | 4181.46 | 606.51 |
| precentral | 15990.94 | 1796.68 | 15929.85 | 1791.05 |
| precuneus | 12865.56 | 1618.69 | 12819.36 | 1616.69 |
| rostralanteriorcingulate | 2963.47 | 479.55 | 2949.78 | 479.97 |
| rostralmiddlefrontal | 21292.13 | 2684.14 | 21165.50 | 2669.35 |
| superiorfrontal | 28758.00 | 3204.70 | 28616.28 | 3197.22 |
| superiorparietal | 17020.90 | 2172.80 | 16961.33 | 2161.06 |
| superiortemporal | 14575.38 | 1645.94 | 14519.78 | 1652.24 |
| supramarginal | 13827.92 | 1891.34 | 13772.95 | 1894.80 |
| frontalpole | 1153.78 | 185.07 | 1150.68 | 186.20 |
| temporalpole | 2478.08 | 309.09 | 2472.20 | 308.04 |
| transverse temporal | 1339.14 | 216.87 | 1333.57 | 217.62 |
| insula | 7586.56 | 857.66 | 7556.20 | 856.70 |
| total | 297024.05 | 28733.94 | 295831.76 | 28686.91 |

## 4.0 Hypothesis Testing and Association Strength

Developing an operational approach to evaluate the meaningfulness of research findings has been a subject of consistent debate throughout the history of statistics^30^. Even with the continued efforts to synthesize systems of statistical inference^31^, the resolution of this issue is unlikely to occur any time soon. Most neuroscientists continue to work within the context of the classical frequentist null-hypothesis significance testing (NHST) paradigm^32,33^, although non-frequentist approaches (e.g. Bayesian, machine learning prediction^34,35^) are increasingly common. Within the NHST framework, researchers attempt to determine which associations are likely “non-null”, or more generally, which associations to prioritize for further examination. For a given dataset, this begins with the choice of a statistical model containing parameters encapsulating the association of interest, and along with a model fitting procedure results in sample estimates of the association parameters. The NHST p-value “…is the probability under a specified statistical model that a statistical summary of the data…would be equal to or more extreme than its observed value”^36^. As indicated in this definition, the p-value depends on the statistical model, with different models potentially giving very different p-values. This underlines the importance of carefully choosing appropriate statistical models and evaluating their assumptions (e.g., models which properly reflect study design elements such as nesting of observations within subjects, subjects within families, and families within sites).

The p-value is distributed over the interval [0,1], uniformly so in the presence of a true null association. Typically, however, a dichotomous decision is reported–should the null hypothesis be rejected? The standard cutoff of *p* ≤ 0.05 is commonly used to guide this decision. The utility of NHST and the arbitrariness of the cutoff value has been debated extensively^36–38^. While we will not relitigate these issues here, we will attempt to address how best to present statistical evidence that leverages the ABCD Study’s large sample size, population sampling frame, and rich longitudinal assessment protocol to enable reliable and valid insights into child and adolescent neurodevelopment. Key takeaways include: 1) the impact of sample size on statistical power and precision of estimates; 2) reporting the magnitude of associations, along with confidence intervals, in addition to p-values; 3) thoughtful control of potentially confounding factors; and 4) ensuring replicable and reproducible results. We cover the first two of these topics in this section, covariate control in Section 5, and briefly touch on replicable and reproducible results, as well as recommendations for statistical analyses and reporting of results in the Supplmentary Materials section.

## 4.1 Power

Statistical power in the NHST framework is defined as the probability of rejecting a false null hypothesis. Power is determined by three factors: 1) the significance level *α*; 2) the magnitude of the population parameter; and 3) the accuracy (precision and bias) of the model estimates. As the p-value is uniformly distributed on the interval [0,1] under the null hypothesis and a well-calibrated statistical model^39^, the significance level *α* is also the Type I error rate, the frequentist probability of rejecting a true null hypothesis. This stands in contrast to the Type II error rate, or the probability of failure to reject a false null hypothesis, denoted by *β* (with power = 1 − *β*). There is always a push-pull relationship regarding the relative seriousness of each error type. Neuroscientists and genomic researchers spend substantial effort attempting to mitigate Type I error rate from high-dimensional data (e.g., via image-wide multiple comparison corrections^40^). Increasing power while maintaining a specified Type I error rate depends largely on obtaining more precise association parameter estimates from improved study designs, more efficient statistical methods, and, importantly, increasing sample size^1,41,42^.

The ABCD Study has a large sample compared to typical neurodevelopmental studies, so much so that one might expect even very small associations to be statistically significant. Possible exceptions to this rule include: 1) analyses of small subgroups; 2) control of many confounding factors and/or complex interactions; 3) rare outcomes; and 4) high-dimensional analyses after multiple testing adjustments. In our experience, not all associations in the ABCD Study are guaranteed to have small p-values even outside of these scenarios. For example, a recent study attempting to replicate the often-cited bilingual executive function advantage failed to find evidence for the advantage in the first data release (NDA 1.0) of the ABCD Study (*n* = 4524)^43^.

Nevertheless, even very small associations are well-powered in the ABCD Study. Figure 3 displays power curves as a function of sample size for different values of absolute Pearson correlations |*r*|. The dashed line in Figure 3 indicates the full ABCD baseline sample size of *n* = 11880. As can be seen, Pearson correlations |r|=0.04 and above have power > 0.99 at *α* = 0.05. Simply rejecting a null hypothesis without reporting on other aspects of the study design and statistical analyses (including discussion of plausible alternative explanatory models and threats to validity), as well as the observed magnitude of associations, is uninformative, perhaps particularly so in the context of very well-powered studies^44^.

**Figure 3:**
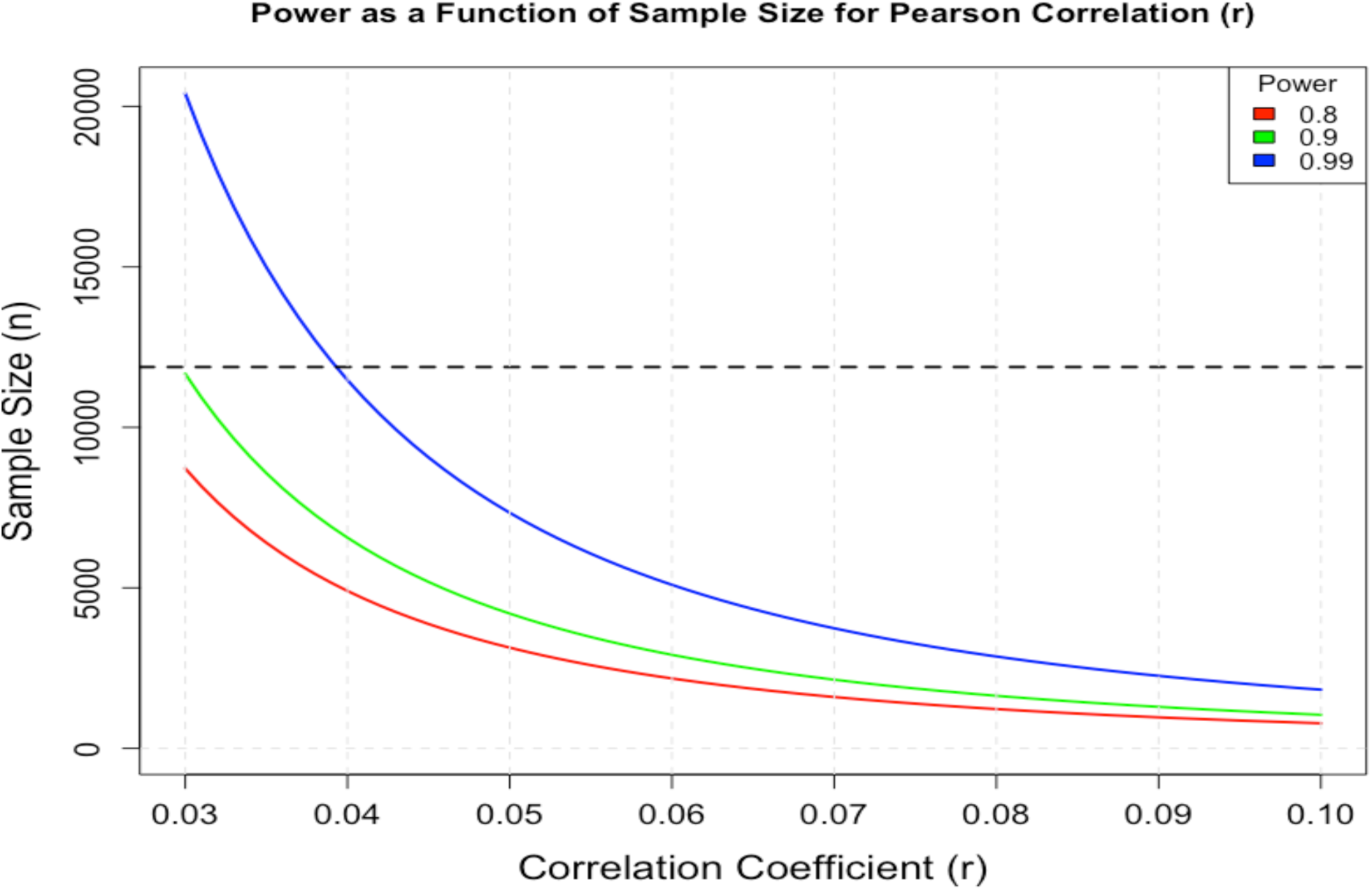
Power vs. Sample Size for Pearson. |*r*|

## 4.2 Precision

The precision of a parameter estimate is its expected closeness to a corresponding population parameter from a given statistical model^45^. Many factors impact precision of parameter estimates, e.g., the magnitude of measurement error and the efficiency of the study design and statistical analysis (Rothman et al.2008, Chs. 10-11)^41^. Crucially, precision is dependent on the sample size *n* — the standard error decreases at the rate of 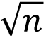 for independent samples. Precision is closely related to power and high levels of precision are especially important to accurately estimate small associations^1^. In fact, underpowered studies can possess a non-negligible probability of obtaining “significant” associations in the wrong direction^46^.

Crucially, increased precision plays an important role in mitigating the impact of publication bias^1^. For example, suppose the strength of an association is quantified by an absolute Pearson correlation |*r*|. Assuming bivariate normality, the interplay of precision and publication bias can be quantified by a simple model involving only the true underlying correlation ρ, the study sample size *n*, and the probability of publication *q_n_*(|*r*|) (e.g., *q_n_*(|*r*|) could be the p-value being below a given threshold; see SM Section S.2).

Figure 4 (left panel) displays this phenomenon in a simulated example of estimated absolute Pearson correlations using bivariate normal samples where the true correlation is *ρ* = 0.10. Five thousand datasets were simulated for each of a range of sample sizes, from *n* = 10 to *n* = 1000. Red lines mark the significance threshold for a Type I error rate of *α* = 0.05, obtained from a normal approximation after a Fisher *z*-transformation utilizing approximate standard errors 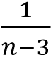. For a sample size of *n* = 10, only 5.8% of samples have an estimated Pearson correlation exceeding this threshold, whereas for *n* = 10000, all estimated correlations exceed the significance threshold in the 5000 simulated datasets. (Note, this essentially recapitulates Figure 3.) The middle panel of Figure 4 displays the expectation of |*r*| vs. n under an extreme selection model whereby only those correlations significant at *α* = 0.05 are published when the true population correlation is *ρ* = 0.10. For *n* = 10, the bias is severe (expectation of 0.71 vs. true value of 0.10), whereas by *n* = 1000 and larger the bias becomes negligible. As a comparison, we display the results of a literature search modified from Feng et al. (2020), which plots 821 brain-symptom absolute correlations derived from 120 publications as a function of study sample size (Figure 4 right panel). The resulting distribution appears qualitatively quite similar to the expectation of |*r*| in the presence of publication bias (middle panel). Thus, *to the extent that publication of results depends on p-values, the bias in the size of published associations will be reduced in larger samples as compared to smaller samples*.

**Figure 4:**
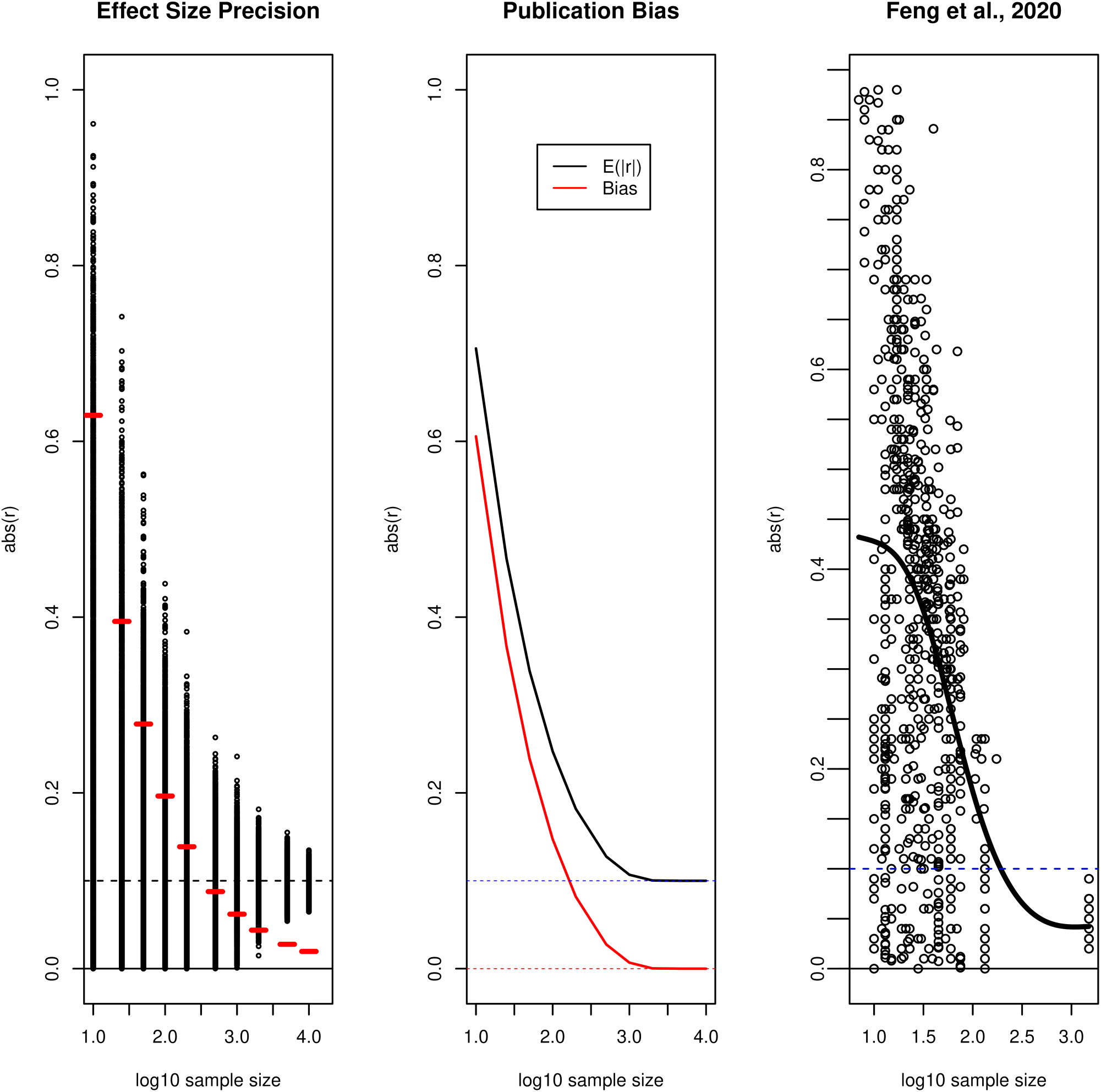
**Sample Size, Reliability, and Publication Bias**

## 4.3 Effect Sizes

An effect size is “…a population parameter (estimated in a sample) encapsulating the practical or clinical importance of a phenomenon under study’’^47^. As most research using the ABCD Study data will not have a direct clinical focus, determining what is meant by “practical importance” will not always be straightforward, as we discuss below. Also note, we are careful to distinguish *effects* (counterfactual, or causal, relationships) from *associations*, which may be impacted by many factors, including selection bias, model misspecification, attenuation due to measurement error, presence of confounders, and/or covariate overcontrol^41,48^. To follow common usage in many treatments on the topic, here we use the term “effect size” rather than “association size,” but we do not intend to imply that unbiased causal effects are necessarily obtainable. We discuss control of confounding factors in the context of the ABCD Study in Section 5.

Effect sizes quantify relationships between two or more variables, e.g., correlation coefficients, proportion of variance explained (*R*^2^), Cohen’s *d*, relative risk, number needed to treat, and so forth^45,49^, with one variable often thought of as independent (exposure) and the other dependent (outcome)^41^. Effect sizes are independent of sample size, e.g., t-tests and p-values are not effect sizes; however, the precision of effect size estimators depend on sample size as described earlier. Consensus best practice recommendations are that effect size point estimates be accompanied by intervals to illustrate the precision of the estimate and the consequent range of plausible values indicated by the data^36^. Table 3 presents a number of commonly used effect size metrics^51,52^. We wish to avoid being overly prescriptive for which of these effect sizes to employ in ABCD applications, as researchers should think carefully about the intended use of their analyses and pick an effect size metric that addresses their particular research question.

**Table 3:** Measures of Effect Size Relevant for ABCD.

|  |
| --- |
| Measures of Strength of Association |
| $r, r_{pb}, r^2, R, R^2, \phi, \eta, \eta^2$<br>Cohen's $f^2$<br>Cramér's $V$<br>Fisher's $Z$ |
| Measures of Strength of Association Relevant for Multiple Regression |
| Standardized regression slope or path coefficient $\beta$<br>Semi-partial correlation $r_{y(x,z)}$ |
| Measures of Effect Size |
| Cohen's $d, f, g, h, q, w$<br>Glass' $g'$<br>Hedges' $g$ |
| Other Measures |
| Odds ratio ($\omega^2$)<br>Relative risk |

## 4.4 Small Effects

As much as the choice of which effect size statistic to report is driven by context, the interpretation of the practical utility of the observed effect size is even more so. While small p-values do not imply that reported effects are inherently substantive, “small” effect sizes might have practical or even clinical significance in the right context^49^.

We may find, as has been true in the majority of published results so far, that most effect sizes reported in analysis of ABCD Study data will be small by traditional standards. Reasons why this may be true include : 1) a broad population-based sample often exhibits smaller effects than narrowly-ascertained clinical samples, perhaps due to ascertainment effects in the latter^4,41,53^; 2) subjects are still young and certain associations, e.g., with psychopathology, may develop more strongly as they progress through adolescence and early adulthood^7^; 3) the large sample size of the ABCD Study increases the power of NHST and the precision of effect size estimates and hence small but non-null effects more easily pass usual significance thresholds compared to estimates from smaller studies.

As described above, known problems of publication bias and incentives for researchers to find significant associations^1,54^ combined with the predominantly small sample sizes of most prior neurodevelopmental studies lead us to expect that true brain-behavior effect sizes are smaller than have been described in the past^55,56^ and attempts to replicate the existing literature using ABCD data will more likely than not result in effect size estimates smaller than prior published effects. Making reference to publication bias and other issues, Ioannidis (2005)^2^ argued that most claimed research findings in the scientific literature are actually false. Although details of the concerns are disputed^57^, some analyses of existing literature provide support for the possibility^58^. We believe a likely scenario is that many published neurodevelopmental associations, while they may not be false positives, do represent severely inflated effect sizes^1,59^.

Reviews of the literature suggest that these issues are pervasive. For example, in a recent metanalysis of 708 individual difference studies in psychology, Gignac and Szodorai (2016)^60^ found that correlations of *r* = 0.11, 0.19, and 0.29 were at the 25th, 50th, and 75th percentiles, respectively. Similarly, in a meta-analysis of mostly treatment/therapy studies, Hemphill (2003)^61^ found that two-thirds of correlations were below *r* = 0.3. According to Cohen’s standards, the majority of studies had reported effect sizes that are below medium, and a good proportion are small (below *r* = 0.10). As such, lack of power due to small effects combined with small samples is a major problem in the field^58^. This is a particularly acute problem for human neuroimaging, where the average power has been estimated to be 0.08, with small-sample studies remaining the current norm rather than the exception^1^. Thus, the extant literature might be represented by effect sizes that are already small, but also inflated relative to the true effect in the population because of the “winners curse”, iterative searching for significant results (“p-hacking”), and publication bias.

In addition to the factors mentioned above, observed effect size estimates may be small for many other reasons, not necessarily related to the magnitude of the underlying mechanistic relationships. These include: 1) measures that may be only weakly correlated with the behavioral and neurobiological constructs of interest^62^; 2) measures with low test-retest reliability and/or high measurement error, which will attenuate effects^63^; 3) measures designed to assess within-person effects, with poor between-person sensitivity^64–66^; and 4) effects that are large within (possibly latent) sub-groups, but which wash-out across the whole sample^67,68^. Many of these factors are germane to some MRI parameters known to have fairly high measurement noise and modest reliability^69–71^, to be susceptible to movement artifacts^72^ (especially in pediatric populations), and to represent only indirect measures of structural and functional indices (e.g., BOLD fMRI measures blood oxygenation and not neuronal activity; diffusion-weighted MRI measures water diffusion and not axon integrity or myelination).

In some contexts (e.g., clinical prediction for individualized treatments) statistically-significant but small effects may not be practically meaningful, and this should be acknowledged. This will likely be the outcome of some proportion of research conducted on the ABCD Study data. The upside of this outcome is that in smaller samples these effects would have ended up in the “file drawer” or estimated with exaggerated magnitude. Thus, the literature will now be able to consider a broader range of results on particular topics of interest, with increased confidence in the likely true size of relationships and with reduced publication biases. The prominent impact of this bias in small-sample research is apparent in the simple simulation presented above but is all but eliminated for large samples, at least when the number of hypothesis tests is not large compared to the sample size.

Finally, we must acknowledge that even if effects are small by usual standards, they should not be inherently dismissed. Small effects may still be important for deciding where to focus attention to understand brain-behavior mechanisms. This has been the case in genomics research where associations of individual loci are tiny for most complex traits but can still be useful for understanding the molecular mechanisms of behavior and identifying potential drug targets for disorders^73^. Moreover, many imperfectly correlated small effects can cumulatively add up to large effects^6,55,56^. Thus, an association can be “practically” important (e.g. useful for informing about brain-behavior mechanisms) even if its effect size is small by traditional standards.

Funder and Ozer (2019)^5^ have recently provided guidelines for reporting effect sizes in terms that are meaningful in context. For example, they argued even small effects (*r* = 0.05) are potentially important if they systematically accrue over time. They reference a classic example of the potential for accumulative consequences of individual behaviors over the long run. In this example, Abelson (1985)^74^ pointed to the correlation between success on a single at-bat in baseball to overall batting average. The effect size is surprisingly small (*r* = .056). However, Abelson argued that systematic differences in single events are nontrivial predictors of future events because the process through which variables operate in the real world is important. Thus, he argued, small effect sizes are meaningful if the degree of potential cumulation is substantial.

In the context of the longitudinal ABCD Study, in which many research questions will be addressed in the context of individual differences, this can be potentially important. As Funder and Ozer point out, “every social encounter, behavior, reaction, and feeling a person has could be considered a psychological ’at bat’” (p. 161)^5^. Effects of this type, which may stem from stable traits of individuals, can have consequences that can add up, and thus small effect sizes, interpreted in the right context, can be meaningful.

## 4.5 Example: Effect Size Estimates

Here we illustrate how the choice of effect size, and the interpretation of its substantive effect, must be made in the context of the research question. For example, Cohen’s *d* and related metrics (see Table 3) assess the magnitude of mean differences between two conditions or groups.But what is not often appreciated is that Cohen’s *d* is insensitive to the proportion of subjects in each group^75^.Thus, Cohen’s *d* might be an appropriate metric for assessing the potential counterfactual impact of an exposure in a given subject (assuming control for confounding factors) but may not be optimal for assessing the public health impact of modifying an existing exposure. Conversely, base-rate-sensitive effect size metrics take into account the difficulty of differentiating phenomena in rare events. If the goal is to assess the impact of an exposure on a population, it is arguable that researchers should opt for an effect size metric that takes the sample base rate into account. For example, the point-biserial correlation *r*_&’_ ^75^ (Table 3) is a similar metric that, unlike *d*, is sensitive to variation in sample base rates.

To illustrate this, we used Cohen’s *d* and point-biserial *r_bs_* to estimate the effect size of a dichotomous “exposure” index: very obese (here defined as a body mass index (BMI) ≥ 30) and a continuous brain “outcome”: restriction spectrum imaging component (N0), a measure sometimes related to cellularity, in the Nucleus Accumbens (NAcc). Recent work has highlighted a potential role of neuroinflammation in the NAcc in animal models of diet-induced obesity^76^. We included baseline data from subjects without missing BMI and NAcc N0 data, also excluding 5 subjects with NAcc N0 values < 0 (leaving *n* = 10659 subjects, of which 184 subjects had BMI ≥ 30, or 1.7%). As can be seen in Figure 5 (upper panels), NAcc N0 values are heavy tailed. We thus use a bootstrap hypothesis testing procedure to obtain quantiles of *d* and *r_bs_*^77^. To account for nesting of subjects within families, at each iteration of the bootstrap one member of each family was first selected at random, and these subjects (along with all singletons) were sampled with replacement 10000 times. Figure 5 (lower panels) presents the bootstrap p-value plots for different null hypotheses^41^. The bootstrap median *d* = 0.801 (95% CI: [0.588,0.907]) and median *r_bs_* = 0.106 [0.081,0.127]. Thus, while in terms of *d* the effect might be considered “large”, *r_bs_* corresponds to a variance explained of roughly 1% and hence would be considered “small” by many researchers.

**Figure 5:**
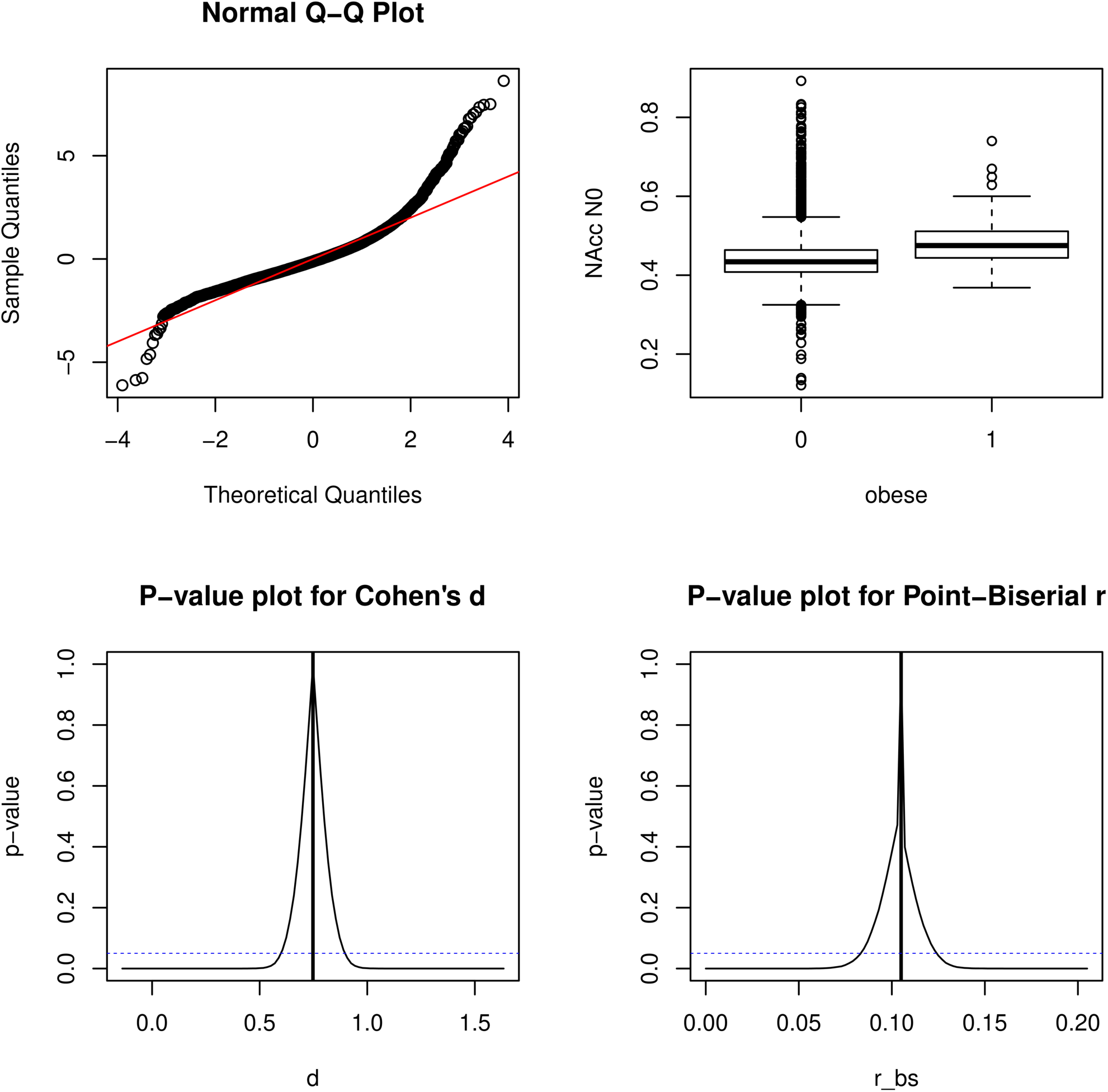
**Association Between Obesity and Nucleus Accumbens RSI N0**

So, what effect size should the researcher report, and which should be emphasized in the interpretation? Our general guidance would be to carefully consider the answer in the context of the research question. Perhaps both could be reported, but if the public health impact of an intervention is considered the *r_bs_* might be more strongly focused on in the discussion of results.

Other factors could affect the calculation of effect sizes. For example, to explore the impact of ABCD sample differences from the ACS data on effect size estimates, we re-ran the analyses using a weighted bootstrap, with probability of sampling proportional to the raked population weights described in Section 3. The weighted bootstrap yielded median *d_wt_* = 0.776 ([0.609,0.951]) and median *r_bs, wt_* = 0.107 ([0.083,0.132]). The median estimates are thus little changed from the unweighted bootstrap medians, though the 95% confidence intervals are wider as expected due to the increased variability in weighted compared to unweighted estimates^26^.

Finally, caution is warranted in interpreting these results as “effect sizes,” as the causal relationship could be from obesity to NAcc N0, from NAcc N0 to obesity, bidirectional, or even non-existent (i.e., due to confounding). We do not adjust for potential confounding factors or their proxies in this example. In light of this, it would be more appropriate to call *d* and *r_bs_*_’_ as computed here “association sizes”. We examine the question of direction of causality using the twin data^78^ in SM Section S.3.

## 5.1 Control of Confounding Variables

Random variation impacts statistical inferences via reduced precision and attenuation of associations. Systematic sources of bias can also threaten the external validity of inferences regarding effects of interest (Rothman et al 2008, Ch. 9^41^). For example, while the ABCD Study endeavored to collect a representative sample of US children born between 2006- 2008, there are small departures from the ACS on some key sociodemographic factors due to self-selection of subjects (Table 1). Using the population weighting described in Section 2, we can adjust the data to more closely resemble that of the ACS in terms of sociodemographic factors assessed in both samples, but this does not guarantee similarity between the ABCD and ACS samples in terms of other variable distributions, if participation in the ABCD Study is related to unobserved factors also related to the variables of interest.

An important challenge to the internal validity of effect estimates from the ABCD Study (and from any observational study) is the likely presence of confounding variables for observed associations. Necessary but not sufficient conditions for a variable to confound an observed association between an independent variable (IV) and a dependent variable (DV) are that the factor is associated with both the exposure and the outcome in the population, but not causally affected by either^79^ (if a variable is causally downstream of the IV or the DV or both, it may be a collider or a mediator^41^). Conditioning on confounders (or their proxies) in regression analyses will tend to reduce bias in effect size estimates, whereas conditioning on colliders or mediators (or their proxies) will tend to increase bias. To make matters more difficult, assessed variables can be proxies for both confounding factors and mediators or colliders simultaneously, in which case it is not clear whether conditioning will improve or worsen bias in effect size estimates. We thus recommend that investigators using ABCD data think carefully about challenges to estimating effects of exposures and perform sensitivity analyses that examine the impact of including/excluding covariates on associations. In the next sections we discuss these topics more thoroughly in the context of conditioning on covariates in regression models.

## 5.1 Covariate Adjustment

Although the inclusion of covariates (sometimes called *control variables*) in statistical models is a widespread practice, determining which covariates to include is necessarily complex and presents an analytical conundrum. The advantages and disadvantages of covariate inclusion in statistical models has been widely debated^80,81^ and reviewed elsewhere^82–84^, so we focus our discussion on the practical implications of covariate adjustment in the ABCD Study data.

Datasets with a rich set of demographic and other variables, like the ABCD Study, lend themselves to the inclusion of any number of covariates. In many respects, this can be seen as a strength of the ABCD Study, but this can also complicate the interpretation of findings when research groups adopt different strategies for what covariates to include in their models. For instance, a recent comprehensive review of neuroimaging studies^85^ found that the number of covariates used in models ranged from 0 to 14, with 37 different sets of covariates across the 68 models reviewed. This review showed that brain-behavior associations varied substantially as a function of which covariates were included in models: some sets of covariates influenced observed associations only a little, whereas others resulted in dramatically different patterns of results compared to models with no covariates. Which variables are appropriately included as confounders in any given analysis depends on the research question, highlighting the need for thoughtful use of covariates.

Covariates are often used in an attempt to yield more “accurate,” or “purified”^84^ estimates of the relationships among the IVs and DV, thereby revealing their “true” associations^82^ (i.e., to eliminate the impact of confounding on observed associations^41^). Under this assumption, the inclusion of covariates implicitly assumes that they are somehow influencing the variables of interest, either contaminating the relationship between the IV and DV or the measurement of the variables of interest. Thus, not controlling for covariates presumably distorts observed associations among the IVs and DV^80,84^. Note that we use “somehow” to emphasize frequent researcher agnosticism regarding the specific role of the covariates included in the model. Because statistical control carries with it major assumptions about the relationships among the observed variables and latent constructs, some of which are generally unspecified and others of which are potentially unknowable, conclusions drawn from models that mis-specify the role of the covariate will be incorrect. When covariates are thought to influence the observed variables of interest but not the latent construct, this is thought of as measurement contamination (Figure 6A).

**Figure 6:**
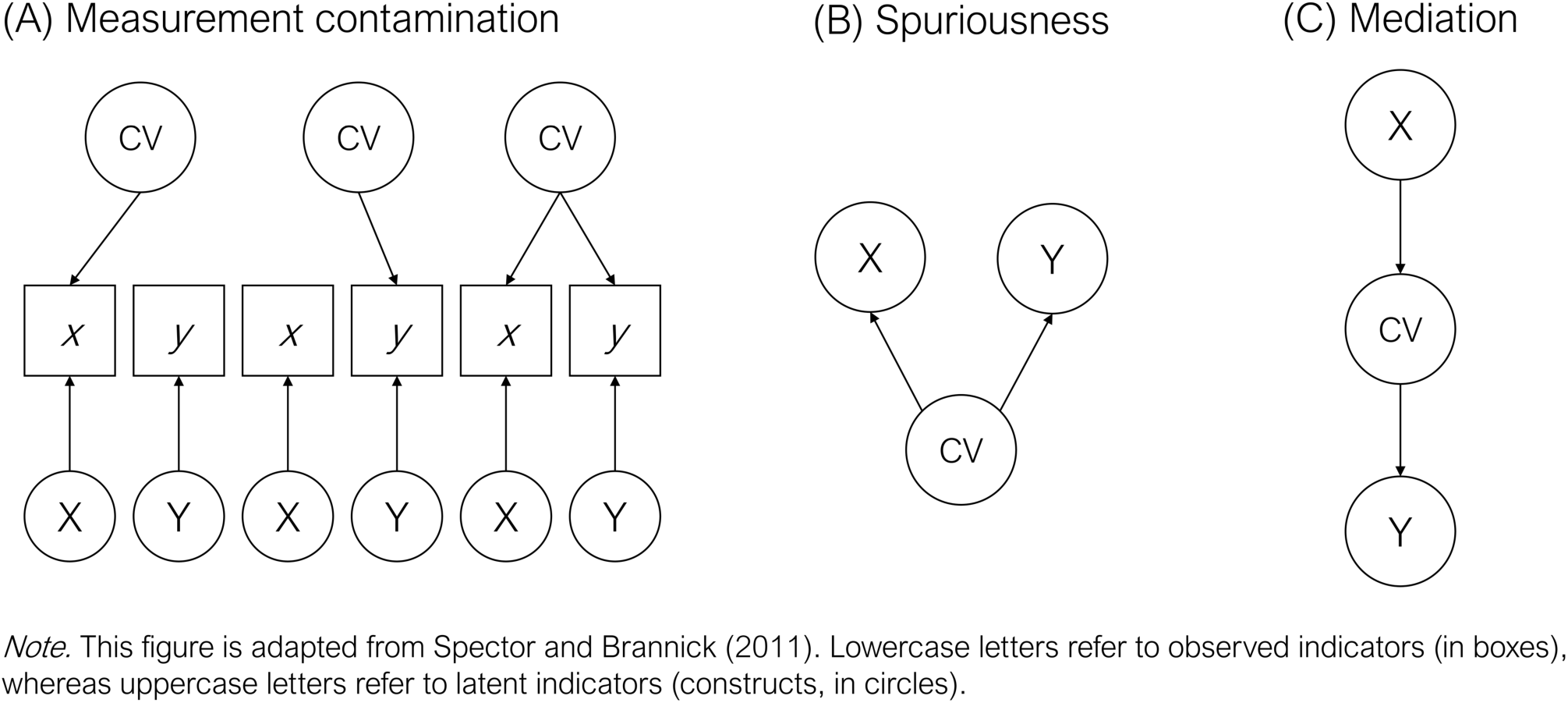
**Models for Measurement Contamination, Spuriousness, and Mediation**

Measurement contamination ostensibly occurs when a covariate influences the observed variables (x and y in Figure 6A). Importantly, a major assumption surrounding the presumption of measurement contamination is that the covariate does not affect the underlying constructs (X and Y in Figure 6A), only their measures. Removing the influence of covariates by controlling for them presumes that absent such control, the association between the IVs and DV is artefactual.

There are also a number of ways in which covariates are thought to influence the latent constructs and not just the measurement of them (see Meehl (1971)^80^ for a thorough discussion). Two such models are spuriousness (Figure 6B) and mediation (Figure 6C). Under a spuriousness (confounding) model, the IV (X) and DV (Y) are not directly causally associated but are both caused by the covariate. Therefore, any observed association between the IV and DV is spurious given that it is caused by the covariate. Under a mediation model, the IV (X) and DV (Y) are statistically associated only through the covariate. Spuriousness and mediation models are generally statistically indistinguishable (though temporal ordering can sometimes assist in appropriate intepretations), and under both models, controlling for the covariate results in a reduced association between the IV and DV. In either case, including covariates can effectively remove effects of interest from the model. At best, this practice obscures rather than purifies relationships among our variables of interest. At worst, this practice can render incorrect interpretations of the true effect. Rather than suggesting that covariates should be avoided altogether, we view them as having an important role in testing competing hypotheses.

In what follows, we offer several general considerations while determining which covariates to use in working with the ABCD data. Afterwards, we provide a worked example focusing on the associations between parental history of alcohol problems and child psychopathology, an important substantive question that has received attention in the literature (e.g., Hesselbrock & Hesselbrock, 1992^86^). We direct interested readers to the following, more thorough treatments of covariate use in statistical modeling^82–84^.

### 5.1.1 Researcher Considerations

*What is the role of the covariate? What is the theoretical model? Could the exclusion and inclusion of the covariate inform the theoretical model?* The practice of simply explicitly specifying the role of the covariate in the model, and even more specifically its hypothesized role in the IV-DV associations, helps avoid including covariates in the model when doing so is poorly justified. Moreover, it encourages thoughtful hypothesis testing. Ideally, explicit justification of the inclusion of each covariate in the model should be included in the reporting of our results. Better yet, as opposed to treating control variables as nuisance variables, a more ideal model would include covariates in hypotheses^83^. As opposed to simply treating an indicator as a covariate whose influence on the IVs and DVs is generally overlooked, we also encourage considering the extent to which the exclusion and inclusion of the covariate could inform the theoretical model. In an explanatory framework, all covariates should be specified *a priori*. In a predictive framework, one can conduct nested cross-validations and model comparisons to find the most robust model with procedurally-selected covariates.

*How do my models differ with and without covariates?* We recommend running models with and without covariates and comparing their results. This practice encourages researchers to better consider the effect of covariates on observed associations. At the same time, engaging in multiple testing can increase Type I error rates. Regarding our suggestion, we encourage a shift away from comparing models on the basis of p-values and instead encourage researchers to compare effect sizes of the predictor of interest in models with and without the covariates. Confidence intervals are critical to compare across models, as the range of plausible effects is more important than the point differences in effect size estimates. The focus on effect sizes as opposed to statistical significance is important given that including many covariates in the statistical model reduces degrees of freedom, in turn increasing standard errors and decreasing statistical power for any given IV. Covariates may be correlated with one another as well, reducing precision and producing large differences in p-values when some variables are included or omitted from a model.

If the effect sizes do not differ as a function of the inclusion of the covariate (e.g., their confidence intervals substantially overlap), one might consider dropping it from the model, but noting this information somewhere in the text. Becker (2005)^87^ offers more suggestions regarding what to do when results from models with and without covariates differ (see also Becker et al. (2016)^83^). Additionally, should one choose to adopt models with covariates included, we recommend placing analyses from models without covariates in an appendix or in the supplemental materials; such a practice will aid in comparison of results across studies, particularly across studies with different sets of covariates in the models.

It is worth formalizing this discussion for situations when there is interest in estimating causal effects: the comparison of potential outcomes, e.g., comparing outcomes for children in ABCD as if all of their parents had alcohol problems, vs. none of their parents having alcohol problems. Two methods that are particularly relevant for estimating causal effects in cohort studies such as ABCD are instrumental variables analyses and propensity score methods. Instrumental variables analyses rely on finding some “instrument” that is plausibly randomly assigned (conditional on covariates), affects the exposure of interest, and is not directly related to outcomes^88,89^.

Here we will focus, though, on propensity score methods as a fairly general purpose tool for estimating causal effects. In general, interpreting a difference in outcomes between exposure groups as a causal effect requires two things: 1) “overlap” (individuals in the two exposure groups are similar to one another on the confounders), and 2) “unconfounded treatment assignment”; that there are no unobserved differences between exposure groups once the groups are equated on the observed characteristics. Propensity score methods^90^ can help assess whether overlap exists, and equate the exposure groups using matching, weighting, or subclassification. Covariates should thus be selected in order to satisfy unconfounded treatment assignment, and as noted above, factors that are “post-treatment” (and thus potentially mediators) should not be included. A benefit of the ABCD Study design is that longitudinal data is available, to measure confounders before exposure and exposure before outcomes, and the large set of potential confounders observed and available to be adjusted for. Sensitivity analyses also exist to assess robustness of effect estimates to a potential unobserved confounder (e.g.,^91^). Finally, methods should be used that account for the probability sample nature of the ABCD cohort, in order to ensure effects are being estimated for the population of interest^93,94^.

### 5.1.2 Example: Covariate Control

Here, we provide a worked example which examines the association between family history of alcohol problems and child externalizing and internalizing psychopathology. The ABCD Study contains a rich assessment of family history of psychiatric problems (e.g., alcohol problems, drug problems, trouble with the law, depression, nerves, visions, suicide) and child psychopathology, including child- and parent-reported dimensional and diagnostic assessments. We will examine the relation between parental history of alcohol problems (four levels: neither parent with alcohol problems, father only, mother only, both parents) and child psychopathology per the parent-reported Child Behavior Checklist (CBCL) Externalizing scores in this example. Based on the earlier-described considerations, we delineate several tiers of covariates to include in the models in sequence (or in a stepwise fashion). The first tier includes “essential” covariates that the researcher views as required to include in the models, the second tier includes “non-essential” covariates, and the third tier includes “substantive” covariates that can inform the robustness of the model, or more generally inform the theoretical model.

Our first tier includes age and sex at birth, which tend to be included in most models. Additionally, the first tier includes a composite of maternal alcohol consumption while pregnant. The inclusion of this covariate is deemed as essential to rule out the possibility that any associations between parental history of alcohol problems and child psychopathology was not due to prenatal alcohol exposure. In this context, maternal alcohol consumption is considered a construct confound. The second-tier covariates include race/ethnicity, household income, parental education, and parental marital status. In the context of this research question, these covariates might be deemed “non-essential” for three reasons. First, there may not be any clear hypotheses surrounding the role of these covariates in the IV-DV associations. Second, there may be reason to think that there are important group differences in the second-tier covariates that are worth exploring and reporting. Third, the researcher might expect that some of the “non-essential” covariates may be causally related to the IVs and DV or may share common causes with them (e.g., they may be proxies for both confounders and mediators or colliders simultaneously). In this example, we do not have specific hypotheses regarding race/ethnicity differences in these associations, but exploratory analyses may be of interest. At the same time, race/ethnicity may be strongly associated with other covariates (e.g., socioecomomic status, adversity), and so researchers must take care when interpreting the impact of its inclusion in the model.

Other “non-essential” covariates including household income, parental education, and parental marital status, may be either causally related to the IVs or DV or may share a common cause. For instance, some data suggest that parental externalizing traits – which are likely to overlap with parental history of alcohol problems – are associated with both increased likelihood of divorce and child externalizing. Importantly, however, parental divorce and child externalizing are not causally related (e.g., Lahey et al., 1998^95^). Similarly, other data suggest that alcohol problems and divorce are genetically correlated^96^. Together these data suggest that demographics may, at least in part, proxy our variables of interest. Here, parental history of alcohol problems may proxy the broader construct of externalizing psychopathology. Moreover, controlling for indicators that share a common cause with our IVs and DVs partials out an important, etiologically relevant part of the phenotype. In doing so, this can obscure true associations between the IV and DV. Based on this information, one might decide to report models with and without these covariates and consider the extent to which differences in these sets of models inform a particular theoretical model.

Figure 7a displays the associations between parental history of alcohol problems and CBCL Externalizing with tier 1 and tier 2 covariates. As you will see, there is a significant linear association between parental history of alcohol problems with tier 1 covariates included, and there is no major difference between the models with and without tier 2 covariates. Given that we deemed tier 2 covariates as “nonessential,” we elected to move forward with tier 1 covariates only. Finally, a third tier of covariates may be used to test the robustness of the associations between parental history of alcohol problems and child psychopathology. We refer to these as “substantive” variables, although the distinction between demographic and “substantive” variables can be arbitrary, like in the case of parental marital status and alcohol problems. As we noted earlier, also available in the ABCD data are parental history of drug use, trouble with the law, and other forms of psychopathology. In Figure 7b, we see that other forms of parental history of psychiatric problems display similar, if not more robust associations, with CBCL Externalizing. Specifically, effects for parental history of other drugs and trouble with the law are significantly more associated with CBCL Externalizing than was parental history of alcohol problems.

**Figure 7:**
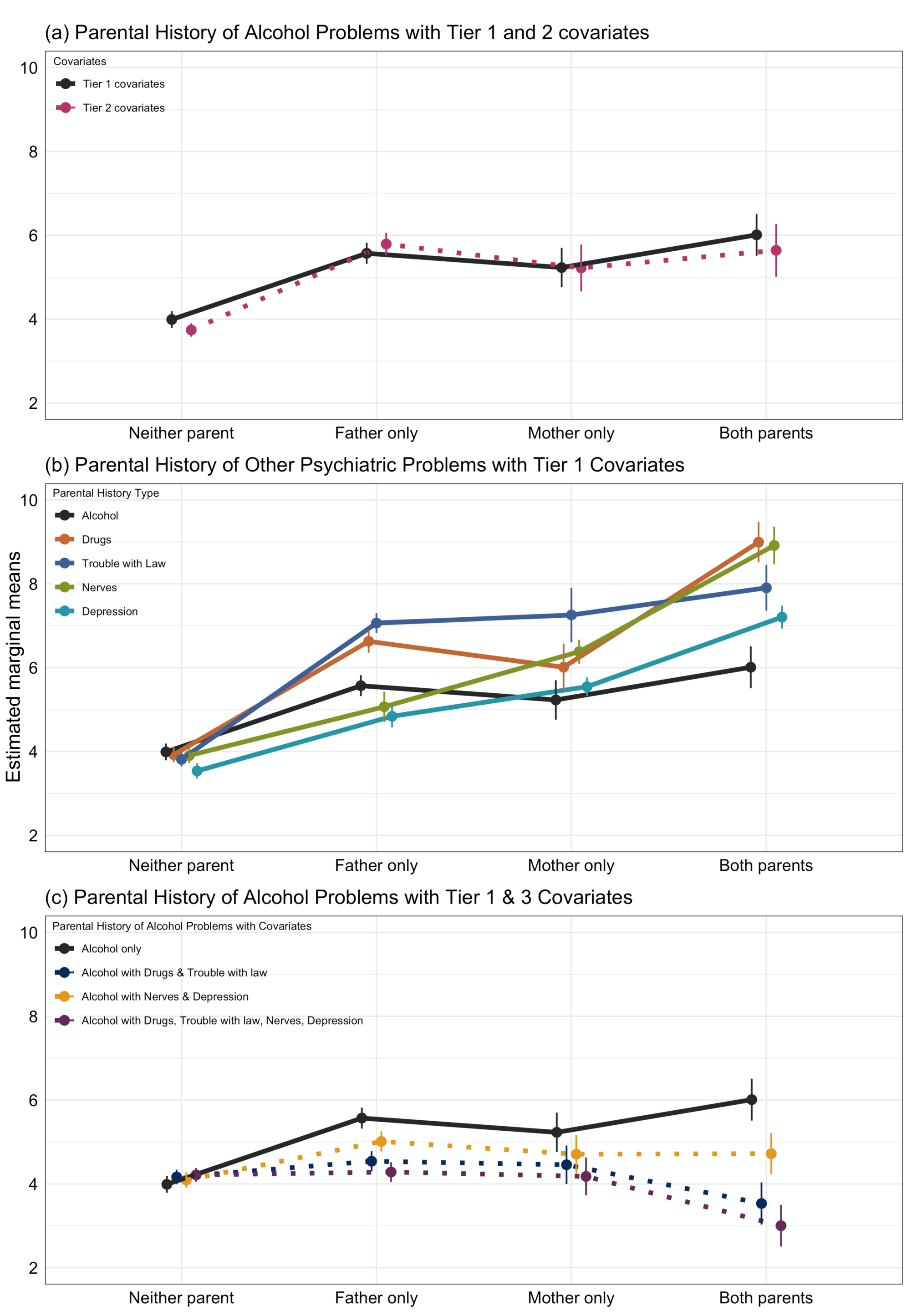
**The association between parental history of alcohol problems and CBCL Externalizing**

Including other forms of externalizing behavior, such as drug use and having trouble with the law, may inform the extent to which the associations between parental history of alcohol problems and child psychopathology are more general to parental history of other externalizing. This seems plausible given research demonstrating significant etiologic (including genetic) associations between numerous forms of externalizing psychopathology (e.g., Kendler et al. (2011)^97^). Figure 7c displays the associations between parental history of alcohol problems and CBCL Externalizing, which became attenuated when parental history of drug problems and trouble with the law were included in the model. This suggests that the associations are general with respect to parental history of externalizing. Because we also saw that other forms of parental history of internalizing problems (i.e., nerves, depression), we can further test whether including them as covariates further attenuates the associations between parental history of alcohol problems and CBCL Externalizing. As we see in Figure 7c, covarying parental history of nerves and depression slightly attenuates the associations between parental history of alcohol problems and CBCL Externalizing, though the effects of covarying parental history of externalizing were stronger.

Both of these tests inform the robustness of the proposed research question. Taken together, we learn from using “substantive” indicators as covariates that the associations between parental history of alcohol problems and CBCL Externalizing may be more general to history of externalizing, or even psychiatric problems more generally. In this case, these “substantive” indicators were not treated as covariates *per se*, but rather variables whose inclusion and exclusion can inform the theoretical model. Determining which covariates should be included in our statistical models is complex and requires considerable thought. We caution against the over-inclusion of covariates in statistical models, and against the assumption that including covariates purifies the associations among our variables of interest; instead their inclusion can obscure rather than purify such associations^98^.

## 5.2 Example: Sensitivity Analysis for Unmeasured Confounding

Unmeasured confounding is a potential threat to internal validity in all observational studies^41^. We present a worked example of how sensitivity analysis can be used with the ABCD dataset to quantify unmeasured confounding. We consider the association between breastfeeding and fluid intelligence, applying the approach of Cinelli and Hazlett (2020)^99^ that allows computation of bias in terms of an unmeasured variable’s association with the outcome and the exposure. The effect size of interest is 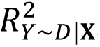, the partial *R*^2^ of the outcome *Y* and the exposure *D* controlling for measured confounds ***X***. An unmeasured confound *Z* is characterized in terms of outcome confounding 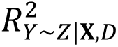, the partial *R*^2^ of the outcome on the unmeasured confound controlling for the measured confounds *X* and exposure *D*, and exposure confounding 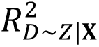, the partial *R*^2^ of the exposure on the unmeasured confound, controlling for measured confounds *X*.

The sensitivity analysis consists of exploring plausible values of 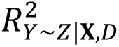 and 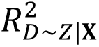 and assessing the impact they would have on the effect strength 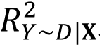, if we did actually correct for *Z*. The primary sensitivity metric recommended by Cinelli and Hazlett (2020)^99^ is the robustness value (*RV_q_* _= 1_), the magnitude of equal outcome and exposure confounding 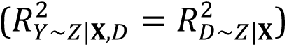 that, after accounting for *Z*, reduces the exposure-outcome association to 0. An additional robustness value, *RV_q_* _= 1, *α* = 0.05_, is similar, but more stringent, and is the equal outcome and exposure confounding needed to merely diminish the effect so that it is no longer statistically significant. The crux of the sensitivity analyses is to establish what are the plausible values of outcome and exposure confounding, 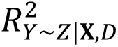, and 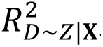 ascertain if they could explain-away the observed effect 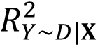.

In this example, the “exposure” is breastfeeding, and the “outcome” is fluid intelligence. A review of the literature finds that maternal IQ is an important covariate but is unavailable in the ABCD dataset. However, we can use related variables in the ABCD dataset to gauge the potential strength of unmeasured confounding and establish whether it is large enough to change our conclusions. We use the **Sensemakr** package, available in R and Stata^100^.

For this analysis, we include only participants that attended the ABCD baseline visit with their biological mother (*n* = 10131). The restriction is important because we want the parental education covariate to reflect the biological mother’s attainment. In addition to maternal education, we select variables that are confounders or strong predictors of neurocognitive performance, and include the ABCD population weights to account for the sampling design^24^. We also included the child’s sex at birth, age at baseline, race/ethnicity, weeks born premature, relationship with biological mother, and school risk and protective factors. We also include the mother’s household income, marital status, tobacco or alcohol use during pregnancy, educational attainment, and age at birth of child. To simplify the analysis, breastfeeding is treated as a binary variable (breastfed, not breastfed). The outcome variable is the NIH Toolbox Fluid Cognition Score, which is a composite of the Flanker, Dimensional Change Card Sort, Picture Sequence Memory, List Sorting, and Pattern Comparison tests^101^.

The results adjusted for all other effects (Table 4) shows that being breastfed is associated with a 1.18-point increase in score compared to children who were not breastfed. The same model reported a strong association between a mother’s education and fluid composite score. We select mother’s education as a variable to benchmark the strength of the unmeasured confounder, suggesting plausible values of 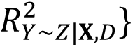 and 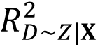.

**Table 4:** Breastfeeding and NIH Toolbox Fluid Composite Score.

|  | NIH TB Fluid Composite Score |  |
| --- | --- | --- |
| | $\beta$ (95% CI) | p-value/t-value |
| Breastfed |  |  |
| No | Reference |  |
| Yes | 1.181 (0.667, 1.696) | $6.89 \times 10^{-6}/4.5$ |
| Maternal Educational Attainment |  |  |
| <HS Diploma | Reference |  |
| HS Diploma/GED | 0.650 (-0.361, 1.662) | 0.207/1.3 |
| Some College | 1.658 (0.754, 2.561) | <0.001/3.6 |
| Bachelor | 3.587 (2.598, 4.576) | $1.25 \times 10^{-12}/7.1$ |
| Post Graduate Degree | 4.088 (3.051, 5.124) | $1.22 \times 10^{-14}/7.7$ |

\* Model adjusted for child sex, age, race/ethnicity, premature birth, relationship with mother, school risk, household income, maternal marital status, tobacco or alcohol use during pregnancy, maternal educational attainment, maternal age at birth of child.

The sensitivity analysis is shown in Table 5. The partial *R*^2^ corresponds to a (*RV*_*q*=1_) of 4.7%, indicating that any unmeasured confounder that explains less than 4.7% of the residual variance in both the treatment and the outcome is not strong enough to fully explain this effect. Considering statistical significance at the *α* = 0.05 level, the *RV_q_* _= 1, *α* = 0.05_ of 2.7% means that the unmeasured confounder needs to explain at least 2.7% of both the treatment and the outcome to make the estimate statistically insignificant.

**Table 5:** Sensitivity Analyses for Breastfeeding and NIH Toolbox Fluid Composite Score.

| Outcome: *NIH Toolbox Fluid Composite Score* |  |  |  |  |  |  |
| --- | --- | --- | --- | --- | --- | --- |
| Treatment: | Est. | S.E. | t-value | $R^2_{Y \sim D \vert \mathbf{X}}$ | $RV_{q=1}$ | $RV_{q=1, \alpha=0.05}$ |
| *Breastfed: Yes* | 1.181 | 0.263 | 4.5 | 0.2% | 4.7% | 2.7% |
| df = 8699 | *Bound (1x mothers education): $R^2_{Y \sim Z \vert \mathbf{X}, D} = 1.3\%$, $R^2_{D \sim Z \vert \mathbf{X}} = 1.8\%$* |  |  |  |  |  |

The role of the benchmark variable is shown in the footer of Table 5, showing that mother’s education has outcome confounding of 1.3% and exposure confounding of 1.8%. Both values are below the RV of 4.7% (and 2.7% for *RV_q_* _= 1, *α* = 0.05_), allowing us to conclude that an unmeasured confounder equal in strength to a mother’s education cannot change our conclusion regarding the effect estimate.

Since the effect of unmeasured confounding depends on two values, 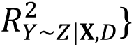 and 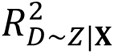, full exploration of confounding requires a plot. Figure 8 shows a t-value contour plot, showing the t-value that would have been observed under different combinations of outcome (y-axis) and exposure (x-axis) confounding, with the critical t-value of 1.98 shown in red. In the lower left, at (0,0), is the original unconfounded result, t-value of 4.5, and two points plotted in red show t-values obtained if an unmeasured confounder had the same (1x) or double (2x) outcome and exposure confounding as the mother’s education benchmark variable. As reflected in Table 5, an unmeasured confounder with characteristics like mother’s education would not eliminate statistical significance, but one with double the confounding effect would change our conclusion. At that point the researcher can discuss the strength of their estimates in a context that has quantified unmeasured confounding. Whether an unmeasured confounder exists that can plausibly change a conclusion depends on domain knowledge and expert judgment. The approach illustrated above allows the researcher to quantify that knowledge and thus measure the impact on effect strength and significance of the signal. We strongly suggest thorough review of the literature prior to selecting a benchmark covariate that has a large but plausible impact on the results.

**Figure 8:**
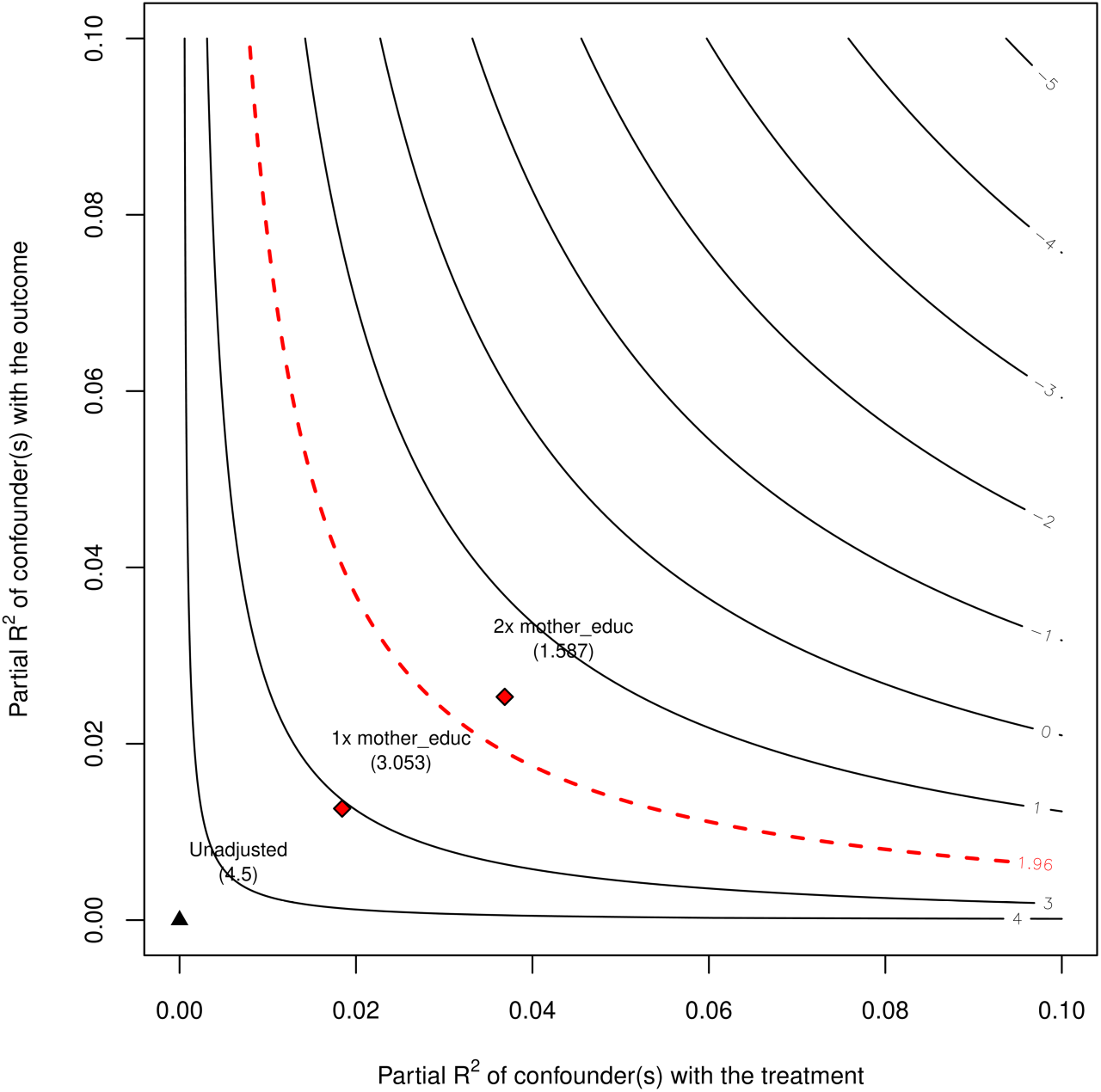
**Plotting Unmeasured Confounding**

## 6.0 Discussion

The sample size of the ABCD Study is large enough to reliably detect and estimate small effect size relationships among a multiplicity of genetic and environmental factors, potential biological mechanisms, and behavioral and health-related trajectories across the course of adolescence. Thus, the ABCD Study will be a crucial resource for avoiding Type I errors (false positive findings) when discovering novel relationships, as well as failures to replicate that result from the replication sample being too small to have sufficient power. Moreover, ABCD will allow for stronger interpretation of non-significant results as they will not be due to low power for all but the tiniest of effect sizes. Other studies in the field suffer from false positives that do not replicate, and overestimation of effect sizes in general, which typically arise from a research environment consisting of many small studies, p-hacking, and publication bias towards positive findings^102^. ABCD will therefore help directly address the replication problems afflicting much of current neuroscience research^1^.

While not of course completely immune to these problems (especially in subgroup and/or high-dimensional analyses), the ABCD Study is much more resistant than are typical small-scale studies, because its large sample size reduces random fluctuations in effect size estimates that occur within small *n* studies. However, with the large number of covariates, high-dimensional space of outcomes and an essentially infinite number of possible modeling strategies, p-hacking and exploitation of random chance remains a possible source of irreproducible results. For example, a recent meta-analysis^103^ found that effects from publications without pre-registration (median *r* = 0.36) skewed larger than effects from publications with pre-registration (median *r* = 0.16). We recommend that researchers consider hypothesis pre-registration (e.g., using the Open Science Foundation framework: https://osf.io/prereg/) and using a registered reports option for publishing results using the ABCD Study data^104^. A template for hypothesis pre-registration for the ABCD Study data can be found in the NDA-hosted ABCD Data Exploration and Analysis Portal (ABCD DEAP, https://deap.nimhda.org/index.php), which is freely accessible to all users with a valid NDA ABCD user ID and password. Over 200 peer-review journals now offer registered reports as a publication format; two of these (*Cerebral Cortex* and *Developmental Cognitive Neuroscience*) have created registered reports options specifically geared for publishing results from the ABCD Study.

Because of the sample size of ABCD, even small effects (e.g., explaining 1% of variation or less) will often be highly significant. In this scenario, it becomes a crucial question how to interpret and utilize the observed relationships and establish their “practical significance.” It is possible that actual (causal) associations found in nature are numerous and small for many outcomes. There is already strong evidence for this possibility: Myer and colleagues (2001)^105^ reviewed 125 meta-analyses in psychology and psychiatry and found that most relationships between clinically important variables are in the r=0.15 to 0.3 range, with many clinically important effects even smaller. Miller et al. (2016)^4^ analyzed associations between multimodal imaging and health-related outcomes in the UKBiobank data. Even the most significant of these explained around 1% of the variance in the outcomes. Thus, like with individual SNPs in a GWAS of complex traits, there are likely many mechanisms involved in producing health outcomes, and each individual observed relationship is a small part of a much larger interacting system.

It is therefore possible that ABCD will predominantly report small effect sizes, simply reflecting the fact that many, if not most, real-world relationships are in fact small. In this scenario, it would be a mistake to dismiss all small effect size relationships for four reasons. First, an ostensibly small effect size might still be of clinical or public health interest depending upon the metric and the importance of the problem^49^. Second, some types of effects (e.g., interactions in field studies) may appear to be small via traditional metrics (e.g., *r*) but represent important, nontrivial effects^106,107^. Third, effects may be small due to imprecise measurement even if the underlying relationships are far from weak. Fourth, even if the effects of individual factors are small, they may cumulatively explain a sizeable proportion of the variation in neurodevelopmental trajectories, a scenario which has recently played out in genome-wide association studies (GWAS) of complex traits^6^. If every small effect were to be thrown away, this would risk never making substantial progress on explaining a substantial amount of total variation in outcomes.

At the same time, it is important that the focus remains on effect sizes, rather than binary “yes or no” assessments of whether data support or reject a particular hypothesis. For example, for the goal of obtaining personally relevant modifiable predictors of substance abuse or other clinical outcomes, prediction accuracy of 75% would correspond to a very-large effect size of around 1.4, accounting for about 30% of the variance. (However, for modifications of variables targeted at a population level or for policy interventions, a smaller effect size might still be important.) Thus, binary judgements on whether associations are “significant” can be fraught with error and give rise to misleading headlines^108^. Worse, Type I or Type II errors (declaring an effect to be significant when it is not real, or absent when it is, respectively) can mislead the field for long periods. Such results could delay the much needed progress in reducing the human and financial costs of mental health and other disorders.

In GWAS, much higher standards of statistical significance are required: typically, one in 20 million rather than the one in twenty value used for single tests. Control of false positive findings in this fashion is essential whenever a very large number of tests are carried out. The neuroimaging data and genomic data being collected in ABCD will be analyzed with the same appropriate adjustments to significance levels when multiple testing is involved. However, there remains a risk that researchers who utilize the public data could fail to observe standard procedures for correcting for multiple testing, not control for design features of the study or measured confounding variables in analyses, or not include effect size estimates in their publications using the ABCD data. Here journal editors and reviewers provide a line of defense against misleading or incorrect reports.

The ABCD Study is collecting longitudinal data on a rich variety of genetic and environmental data, biological samples, markers of brain development, substance use, and mental and physical health, enabling the construction of realistically complex etiological models incorporating factors from many domains simultaneously. While establishing reproducible relationships between pairs (or small collections of measures) in a limited set of domains will still be important, it will be crucial to develop more complex models from these building blocks to explain enough variation in outcomes to reach a more complete understanding or to obtain clinically-useful individual predictions. Multidimensional statistical models must then incorporate knowledge from a diverse array of domains (e.g., genetics and epigenetics, environmental factors, policy environment, ecological momentary assessment, school-based assessments, and so forth) with brain imaging and other biologically-based measures, behavior, psychopathology, and physical health, and do this in a longitudinal context. The sample size, population nature, duration of study, and, importantly, the richness of data collected in ABCD will be important for attaining this goal.

## Acknowledgments

We thank the families who have participated in this research. We also thank the ABCD Biostatistics Work Group. The corresponding author was supported by United States National Institutes of Health, National Institute on Drug Abuse: 1U24DA041123-01 (Dale).

Data used in the preparation of this article were obtained from the Adolescent Brain Cognitive Development^SM^ (ABCD) Study (https://abcdstudy.org), held in the NIMH Data Archive (NDA). This is a multisite, longitudinal study designed to recruit more than 10,000 children age 9-10 and follow them over 10 years into early adulthood. The ABCD Study® is supported by the National Institutes of Health and additional federal partners under award numbers U01DA041048, U01DA050989, U01DA051016, U01DA041022, U01DA051018, U01DA051037, U01DA050987, U01DA041174, U01DA041106, U01DA041117, U01DA041028, U01DA041134, U01DA050988, U01DA051039, U01DA041156, U01DA041025, U01DA041120, U01DA051038, U01DA041148, U01DA041093, U01DA041089, U24DA041123, U24DA041147. A full list of supporters is available at https://abcdstudy.org/federal-partners.html. A listing of participating sites and a complete listing of the study investigators can be found at https://abcdstudy.org/consortium_members/. ABCD consortium investigators designed and implemented the study and/or provided data but did not necessarily participate in analysis or writing of this report. This manuscript reflects the views of the authors and may not reflect the opinions or views of the NIH or ABCD consortium investigators. The ABCD data repository grows and changes over time. The ABCD data used in this report came from NIMH Data Archive Release 2.0.1 (DOI 10.15154/1506087). DOIs can be found at https://nda.nih.gov/abcd.

## Supplementary Materials

### S.1 ABCD Study Aims

The major aims of the ABCD Study include:

• **Aim 1**: Development of national standards of healthy brain development;
• **Aim 2**: Description of individual developmental trajectories in terms of neural, cognitive, emotional, and academic functioning, and influencing factors;
• **Aim 3**: Investigation of the roles and interaction of genes and the environment on development;
• **Aim 4**: Examination how physical activity, sleep, screen time, sports injuries (including traumatic brain injuries), and other experiences affect brain development;
• **Aim 5**: Determination and replication of factors that influence the onset, course, and severity of mental illnesses;
• **Aim 6**: Characterization of the relationship between mental health and substance use;
• **Aim 7**: Specification of how use of different substances affects developmental outcomes, and how neural, cognitive, emotional, and environmental factors influence substance use risk, involvement, and progression.

### S.2 Effects of Publication Bias

Let (*X*, *Y*) denote random variables with population correlation *ρ* and let 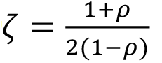 denote the Fisher z-transformation of *ρ*. Further, let *r_n_* denote the Pearson correlation based on a sample of size *n* independent draws of (*X*, *Y*) and 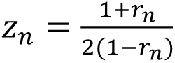 is its Fisher z-transformation. It is well known that *z_n_* is approximately normally distributed with mean *ζ* and standard error 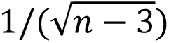.^109^ Finally, let *q_n_*(|*r_n_* |) denote the probabilty that a given *r_n_* is published, dependent only on the sample size *n* and the absolute value of the observed correlation, |*r_n_* |. For example, if signficance at the *α* = 0.05 level increases publication probability, then 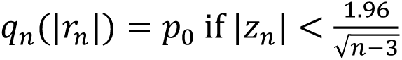 and *q* (|*r_n_* |) = *p*_1_ otherwise, where 0 ≤ *p*_0_ < *p*_1_ ≤ 1. As an extreme case, *p*_0_ = 0 implies only “signficant’’ results are published. More generally, we assume 0 ≤ *q_n_*(|*r_n_* |) ≤ 1 for all *n* and |*r*_n_ | and that the set *S* = {*r_n_* |*q_n_*(*r_n_*) > 0} has positive Lebesgue measure. Given the above model, the probability density function of |*z*_n_| is given by 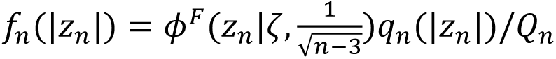, where *ϕ^F^* is a folded normal density and the support of *f_n_* is on the non-negative real line. *Q_n_* is a normalizing factor given by 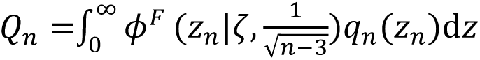. Letting ℎ denote the inverse Fisher z-transformation, the expectation of |*r_n_* | under the publication bias model is then given by 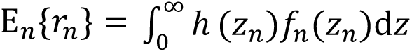. Code for computing the expected value and bias of |*r_n_*| as an estimator of *ρ* is given in the ABCD Biostatistics R package at https://github.com/ABCD-STUDY/ABCD-BIOSTATISTICS/.

### S.3 Example: Direction of Causation Model for BMI and NAcc N0

Multilevel twin models were used to assess the relationship of body mass index (BMI, labeled A in SM Figure 1), with Restriction Spectrum Imaging (RSI) Nucleus Accumbens N0 component (NAcc N0, labeled B in SM Figure 1). We used the Direction of Causation (DoC) model^78^ on all same-sex twins with known zygosity and no missing BMI or NAcc N0 data () in the baseline data (NDA Release 2.0.1). DoC models are Structural Equation Models (SEMs). DoC models exploit the fact that the implied covariance structure of cross-twin cross-trait bivariate data differs based on the causal direction of (A to B, B to A, or bidirectional) under the assumption that the unique and common components of environmental effects are independent between the two variables. DoC models were fit using the R package **OpenMx**^110^.

A to B, B to A, and bidirectional p-values were 0.606, < 0.00001, 0.810, respectively. Small p-values imply the specified model has significantly worse model fit compared to the fully saturated model. Thus, B to A fit substantially worse than the saturated model, whereas A to B and bidirectional models fit similarly to the fully saturated model. The BMI to NAcc N0 standardized effect was 0.203 (95% CI: [0.177,0.229]), implying that a one standard deviation increase in BMI leads to 0.203 standard deviation increase in NAcc N0. Note, these results do not provide evidence for causal relationship *per se*, but rather that, if the assumptions of the DoC model are true, the direction of BMI to NAcc N0 is much more supported by the data than *vice versa*.

### S.4 Reproducible Research

The cornerstone of reproducible research is transparency. Transparency is achieved through annotating and sharing (with others through ‘publication’) precisely ‘what operations’ were performed on ‘what data’ in a fashions that could potentially be re-executed by someone in an attempt to replicate, and ultimately document the generalizability of, the original finding. Deconstructing this statement results in “annotation of the operations” and “annotations of the data” in a way that appropriately authorized ‘others’ can repeat.

#### Annotation of the Data

Raw data comes from multiple sources, including MRI scanners and clinical and behavioral assessments. Annotation of the data sources involves making sure that the data files, as carried forward into analysis or publication, retain a description that is completely self-describing. Self-describing data is necessary to maximize the utility of the data to others, and to minimize the burden on the collector for supporting the future uses of that data. Data that is acquired completely electronically (MRI scans, for example) are already accompanied by comprehensive documentation of the complete set of acquisition descriptions in the scanner-generated DICOM data files. Data that is captured from more locally-generated assessment frameworks (RedCap data entry, pen and pencil forms, etc.) require the investigator to add the appropriate annotation to the local database of data files. This annotation needs o encode precisely what form (and version) is being used and the semantic meaning of the measure (to facilitate interoperation with similar, but not identical measures from other datasets). In large data collections, such as ABCD, this can be a very daunting task, but not doing it limits the utility of hte data going forward, or results in even more time consuming addition of such annotation at a future date. Libraries of annotated markup for reuse in individual labs and studies that lower the barrier to generation of annotated data, and facilitates the annotation of the differences between a local data collection and other similar collections are becoming available (see, for example, ReproSchema, https://github.com/ReproNim/reproschema).

#### Annotation of the Operations

Annotations of the operations that are performed on the data to generate results include data processing and statistical assessments. Such description includes what operations were performed and what computational environment performed the operations. While indications of what specific software tools and versions of the tools were used are a good start, a full descriptions requires indication of he complete parameter set used, and the details of the processing approach (data analysis script) that was employed in order to document the analysis process completely. Similarly, simply stating what operating system and hardware was used does not completely specify the execution environment sufficiently to enable re-execution of the process since details of operating system version, libraries, environment variables, etc. can all impact the details of software results (See Glatard, et al., citation). For this reason, use of virtualized or containerized environments that both completely specify the hardware configuration and do so in a easily sharable and re-deployable fashion is highly recommended.

Finally, to facilitate accessibility to others, all elements of an analysis should be ‘published’. Publication includes publication in the formal traditional peer-reviewed sense, but can also include non-peer-reviewed ‘self’ publication through sharing to publicly accessible resources. Both the data (including initial raw data and the complete results of the analysis), and the operations that were applied to that data (processing scripts and the computational environment, etc.) needs to be accessible to others in order to confirm and generalize a given finding. In summary, given these objectives, a number of themes pervade these best practices. These include:

• Publish everything (including raw data, annualized derived results, processing workflows, etc.) - so that others can access;
• Version control everything - so that you know what you did, and when and why you changed what you did;
• Annotate everything so that you can others can understand your data and results and re-use more easily;
• Use standards - so that others can access what you’ve done more easily; and
• Use containers - so that others can re-do what you did.

### S.5 Recommendations for Analytic Procedures and Reporting of Results

#### Analytic Procedures

• Use analytical methods appropriate for the study design (e.g., mixed models to account for nesting within families)
• Check whether model assumptions hold
• Perform sensitivity analyses of the impact of different model choices and modeling assumptions
• Assess the impact of models with and without (sets of) covariates
• Use appropriate models for a given outcome distribution
• Perform model fit comparisons for competing, equally-plausible models
• Differentiate genuine hypothesis testing from exploratory analyses
• Adjust for multiple testing when appropriate
• Estimate associations robust to overfitting (e.g., using K-fold cross-validation)

#### Reporting Results

• Don’t just report p-values, also report effect sizes (with confidence intervals)
• Choose effect sizes that make sense for what you are attempting to demonstrate
• Report the number of tests you have done
• Show all analyses (even if they end up in the Supplementary Materials)^111^
• Avoiding using causal language, explicitly or implicitly
• Display actual data along with model fits when possible
• Try to provide enough information for others to use results in meta-analyses (e.g., PRISMA and MOOSE guidelines)
• Acknowledge when alternative models with different interpretations could fit the data equally well
• Share your analysis scripts with others - analyses of ABCD Study data should be completely reproducible for others with valid access to the data
• Adhere to reporting standards for observational studies – STROBE guidelines: https://www.equator-network.org/reporting-guidelines/strobe/ – MOOSE guidelines: https://www.elsevier.com/ data/promis_misc/ISSM_MOOSE_Checklist.pdf – Additional reading on best practices for reporting results from observational studies112–115

**SM Table 1:**
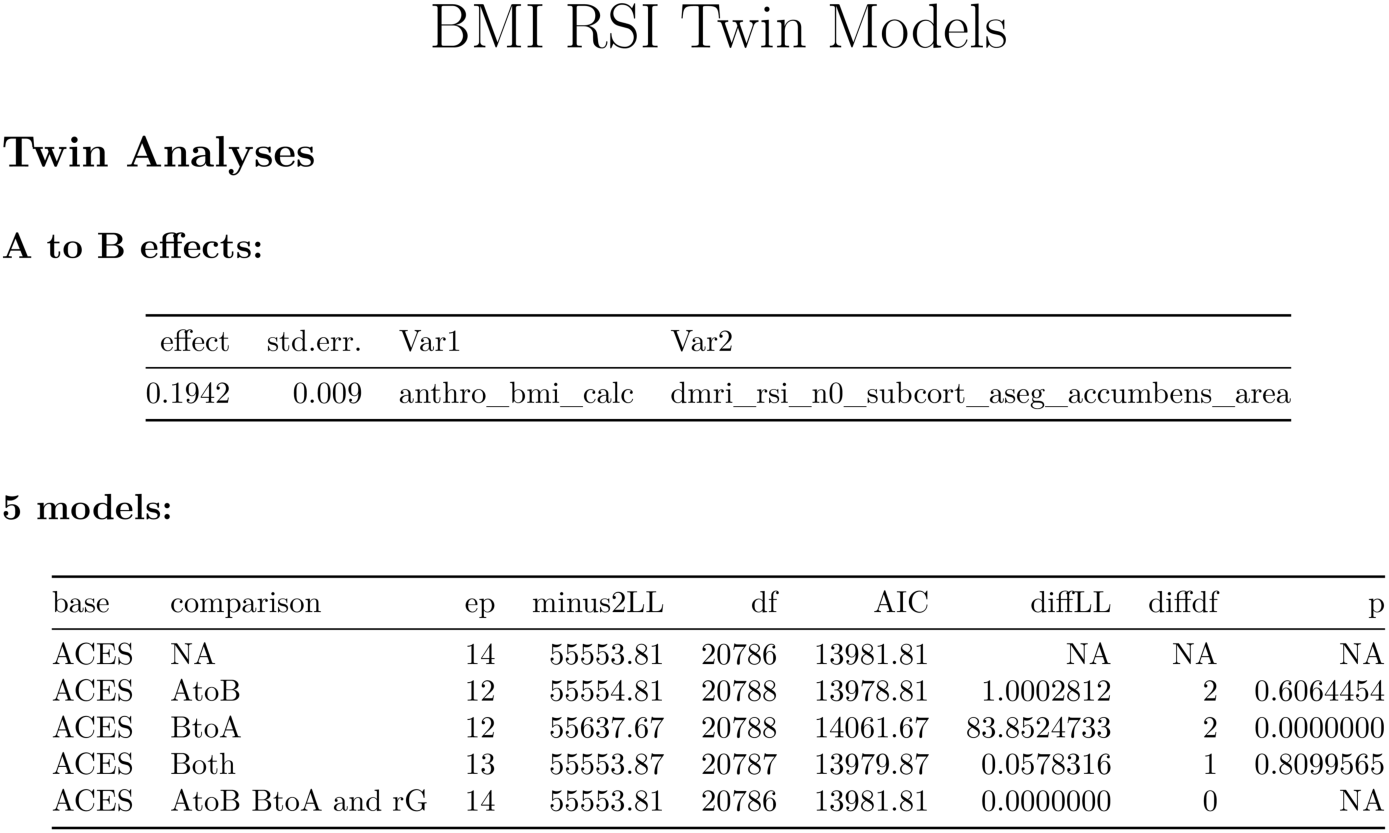
Direction of Causation Model for BMI (A) and NAcc N0 Component (B)

